# Structural insight into the formation of lipoprotein-β-barrel complexes by the β-barrel assembly machinery

**DOI:** 10.1101/823146

**Authors:** Juliette Létoquart, Raquel Rodriguez-Alonso, Van Son Nguyen, Guennaëlle Louis, Antonio N. Calabrese, Sheena E. Radford, Seung-Hyun Cho, Han Remaut, Jean-François Collet

**Author notes:** co-first authors (Juliette Létoquart, Raquel Rodriguez-Alonso).

## Abstract

The β-barrel assembly machinery (BAM) inserts outer membrane β-barrel proteins (OMPs) in the outer membrane of Gram-negative bacteria. In Enterobacteriacea, BAM also mediates export of the stress sensor lipoprotein RcsF to the cell surface by assembling RcsF-OMP complexes. Although BAM has been the focus of intense research due to its essential activity in generating and maintaining the outer membrane, how it functions remains poorly understood. Here, we report the crystal structure of the key BAM component BamA in complex with RcsF. BamA adopts an inward-open conformation, with the lateral gate to the membrane closed. The globular domain of RcsF is lodged deep inside the lumen of the BamA barrel, binding regions proposed to undergo an outward and lateral opening during OMP insertion. Our structural and biochemical data indicate a push-and-pull mechanism for RcsF export upon conformational cycling of BamA and provide a mechanistic explanation for how RcsF uses its interaction with BamA to detect envelope stress. Our data also suggest that the flux of incoming OMP substrates is involved in the control of BAM activity. Overall, the structural insights gleaned here elucidate a fundamental biological process and suggest a new avenue for antibiotic development.

---

The vast majority of proteins inserted in the outer membrane of Gram-negative bacteria adopt a β-barrel conformation. Their assembly depends on the activity of the conserved β-barrel assembly machinery (BAM), whose core component is the OMP85-family protein BamA ^1,2^. BamA is an outer membrane 16-stranded β-barrel with a large periplasmic extension consisting of five POlypeptide TRansport-Associated (POTRA) domains at its N-terminus ^1^. Structures of BAM have shown that BamA can adopt two conformations: an outward-open conformation ^3-5^, in which the β-barrel domain opens between strands β1 and β16 to open a lateral gate to the membrane, and an inward-open conformation ^5,6^, in which the lateral gate is sealed while a periplasmic entry pore to the barrel lumen is open. In the bacterium *Escherichia coli*, four accessory lipoproteins (BamB, BamC, BamD, and BamE) complete BAM, forming a pentameric holocomplex ^7,8^. BamBCDE are anchored to the outer membrane by a lipid moiety but reside in the periplasm. BamB and BamD directly bind the POTRA domains of BamA, while BamC and BamE bind BamD ^1,2^. Although all components are required for efficient assembly of *E. coli*’s diverse set of OMPs, only BamA and BamD are essential and conserved throughout Gram-negative bacteria ^1,2^. Despite important structural and functional insights during 15 years of intense scrutiny, crucial questions remain unsolved regarding BAM’s mechanism. In particular, the functional importance of BamA cycling between the outward-open and inward-open conformations remains unclear, as are the respective contributions of the various BAM components to OMP assembly ^9^.

The primary function of BAM is the assembly of OMPs and, when necessary, the translocation of their associated extracellular domains across the outer membrane. More recently, BAM has also been implicated in export of the outer membrane lipoprotein RcsF to the cell surface ^10,11^ via the assembly of complexes between this lipoprotein and three abundant OMPs (OmpA, OmpC, and OmpF) ^10,11^. Support for the involvement of BAM in RcsF export comes from *in vivo* crosslinking experiments in which a complex between RcsF and BamA, considered to be an intermediate in the formation of RcsF-OmpA/C/F complexes, was trapped ^10,11^. Further, in cells lacking BamB and BamE, RcsF accumulates on BamA and causes a lethal block to BAM-mediated OMP assembly, suggesting that OMPs and surface-exposed RcsF exploit at least partially overlapping assembly routes ^12,13^.

RcsF functions as an envelope stress sensor capable of mounting a protective response when damage occurs in the peptidoglycan or in the outer membrane ^14,15^. Interestingly, we previously determined that sending RcsF to the surface is part of a cellular strategy that enables RcsF to detect damage in the cell envelope. Under stress conditions, newly synthesized RcsF molecules fail to interact with BamA ^10^: they are not exported to the surface and remain exposed to the periplasm, which allows them to trigger the Rcs signaling cascade by reaching the downstream Rcs partner in the inner membrane ^16^. Thus, surface exposure is intimately linked to the function of RcsF. However, the molecular details of the BamA-RcsF interaction, how BAM orchestrates the export of RcsF with OMP assembly, and what prevents RcsF from interacting with BamA under stress conditions remain unknown.

Here we sought to address these questions by obtaining structural information about the interaction between BamA and RcsF. In a series of exploratory experiments, we co-overexpressed RcsF with the BamAB sub-complex, or with the BamABCDE holocomplex; both BamAB-RcsF and BamABCDE-RcsF could be detergent-extracted from the membrane and purified via affinity chromatography using a His-tag on the N-terminus of BamA (Fig. 1). Using native gel electrophoresis, we confirmed that RcsF binds BamABCDE, and not only BamAB (Extended Data Fig. 1a, b, c). However, whereas BamAB-RcsF was stable and could be purified to homogeneity by size-exclusion chromatography, BamABCDE-RcsF was unstable (Extended Data Fig. 1d). The BamAB-RcsF complex was crystallized and its structure solved to 3.8 Å resolution by molecular replacement using the structures of BamA and RcsF (PDB: 5D0O and 2Y1B, respectively; Extended Data Table 1). While this structure contained BamA and RcsF (Fig. 2), BamB was absent, perhaps as a result of proteolytic degradation, as reported previously for this protein when crystallization of the *E. coli* BAM complex was attempted ^4,5^. The asymmetric unit contained two BamA-RcsF conformers, although for one of them, no unambiguous electron density was observed for POTRA domains 1, 2, 3, and 5 (Extended Data Fig. 2a, b). In both BamA copies, RcsF was lodged deep inside the lumen of the BamA β-barrel (Fig. 2a; Extended Data Fig. 2c). The latter was found in an inward-open conformation closely matching that reported in structures of *E. coli* BamABCDE (^6^, with a root mean square deviation of 0.9 Å for 383 equivalent C*α* atoms in the BamA β-barrel of PDB: 5D0O) or BamA truncates lacking POTRA domains 1-4 or 1-5 ^17-20^. Inside BamA, RcsF contacts two BamA loops protruding into the β-barrel: (1) extracellular loop L6 (^e^L6; ∼77 Å^2^ buried surface area, one putative H bond; note that at 3.8 Å resolution, amino-acid sidechain positions cannot be unambiguously determined), and (2) the periplasmic loop connecting strands 7 and 8 (^p^L4; ∼140 Å^2^ buried surface area, one putative H bond) (Fig. 2, 3a). Although contacting RcsF, these loops retain a conformation closely matching that seen in inward-open apo BamA structures (Fig. 3b). However, the main BamA-RcsF contact occurs through the luminal wall of the BamA β-barrel, encompassing ∼1100 Å^2^ of buried surface area and comprising up to 15 putative H-bonds (Fig. 2). This RcsF-BamA β-barrel interaction can be divided into three zones. Zone 1 (Z1) consists of perhaps nine H bonds formed by BamA residue 488 and residues 463, 465, and 466 in the loop connecting β3 and β4 and contacting the RcsF loop connecting β1 and α1 (L^β1-α1^) (Fig. 2b, c, Fig. 3a, c). Zone 2 (Z2) is made of perhaps four H bonds formed by BamA residues 592 and 634, located above ^p^L4 (Fig. 2c, 3a).β16, one of the components of the proposed lateral gate of the BamA β-barrel, constitutes the third zone (Fig. 2b, c, 3a). The bottom of RcsF protrudes out of the BamA β-barrel into the periplasm, residing in close proximity to POTRA domains 3-5 (Fig. 2a, b). As a result, RcsF sterically pushes POTRA5 outward, causing a 26° rotation compared to the inward-open conformation found in apo BamA structures ^5,6^ (Fig. 3b). Although the lipid anchor of RcsF and the N-terminal disordered linker (residues 16-50) ^21,22^ are not apparent in this structure, the position of RcsF is compatible with the lipid anchor residing in the inner leaflet of the outer membrane. Of note, the binding interface between RcsF and BamA does not overlap with the binding sites of BamA for its accessory lipoproteins (Extended Data Fig. 3). Consistent with this observation, the RcsF-BamA interaction is compatible with the binding of BamBCDE, as determined experimentally (Fig. 1; Extended Data Fig. 1).

**Figure 1.**
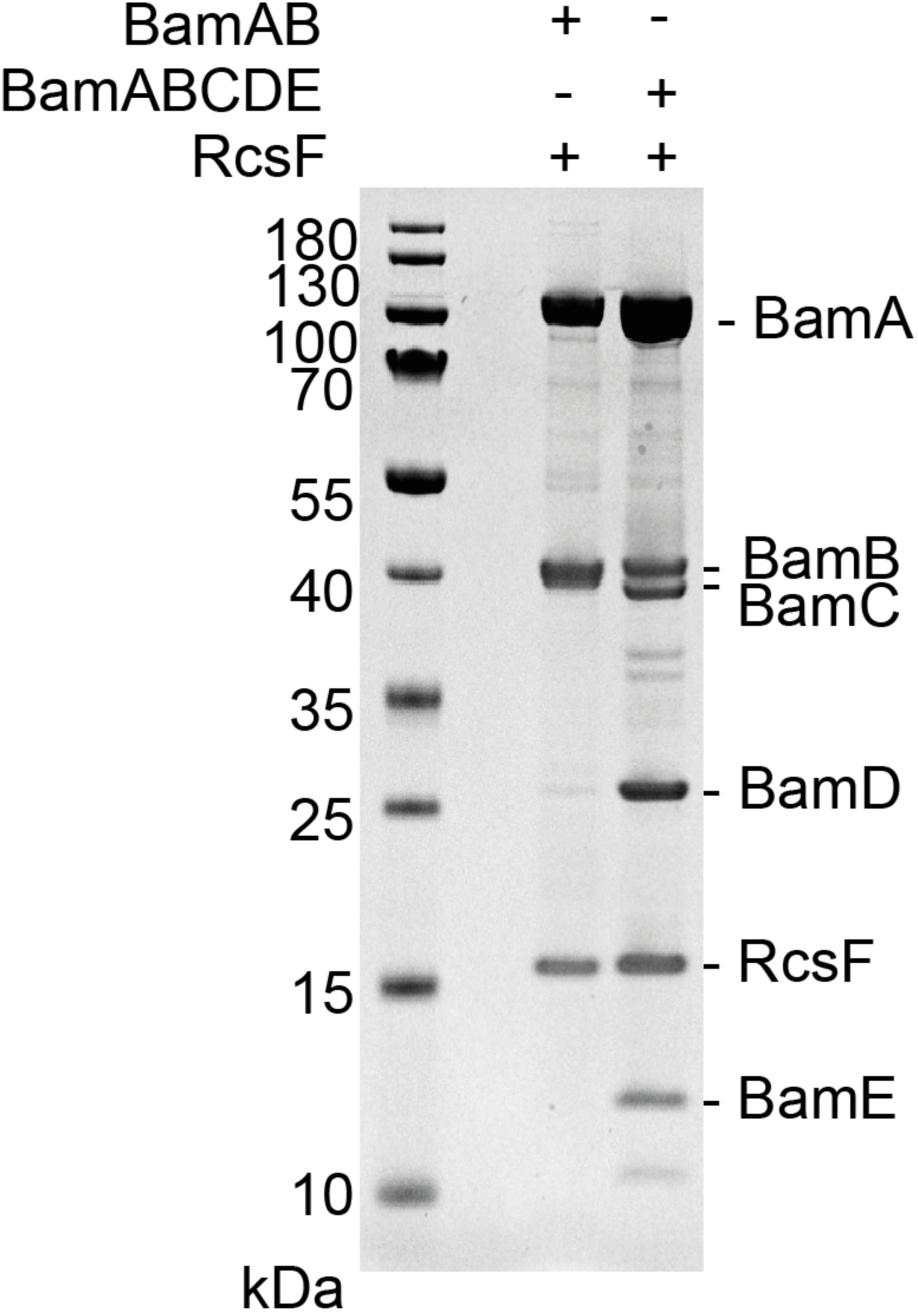
RcsF forms a complex with BamAB and BamABCDE. SDS-PAGE of purified BamAB-RcsF and BamABCDE-RcsF. Plasmids pJH118 and pRRA1 were each co-expressed with pSC216 (RcsF) to yield BamAB-RcsF and BamABCDE-RcsF, respectively.

**Figure 2.**
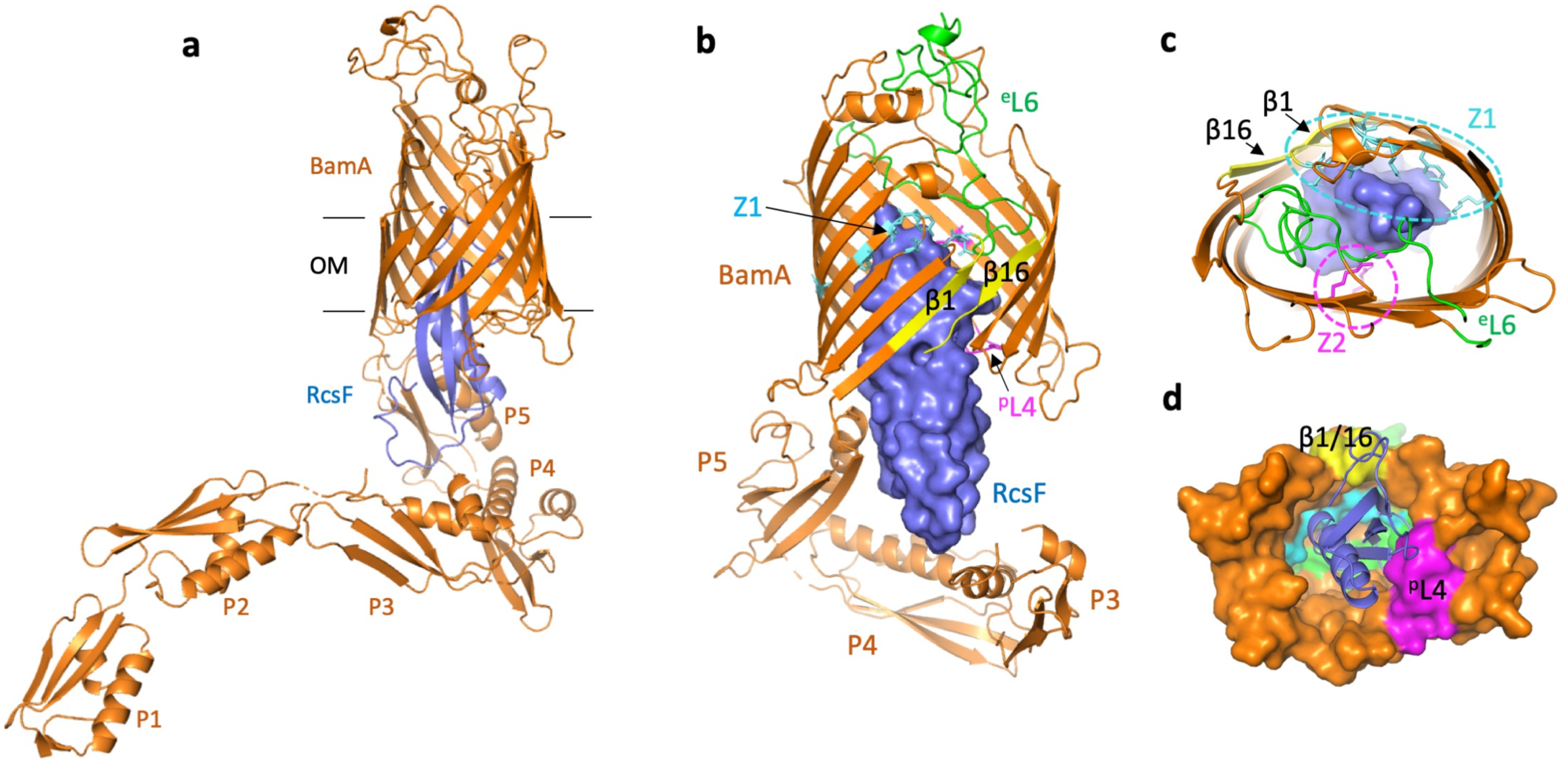
Structure of the BamA-RcsF complex. **(a)** Ribbon diagram of the BamA-RcsF complex in side view. BamA, gold; RcsF, blue. **(b, c)** Front (b) and extracellular (c) views of BamA-RcsF, with RcsF shown as a solvent-accessible surface. POTRA domains 1 and 2 have been omitted for clarity. BamA ^e^L6, green; ^p^L4, magenta. Putative RcsF-interacting residues in contact zones Z1 and Z2 of the BamA β-barrel are colored cyan and magenta, respectively, and shown as sticks. Strands β1 and β16, which form the proposed “lateral gate” of the BamA β-barrel ^1^, are yellow. **(d)** Periplasmic view of the BamA-RcsF complex, with the BamA β-barrel shown as a solvent-accessible surface and RcsF as a ribbon. Colors are as in panels b and c. POTRA domains were omitted for clarity.

**Figure 3.**
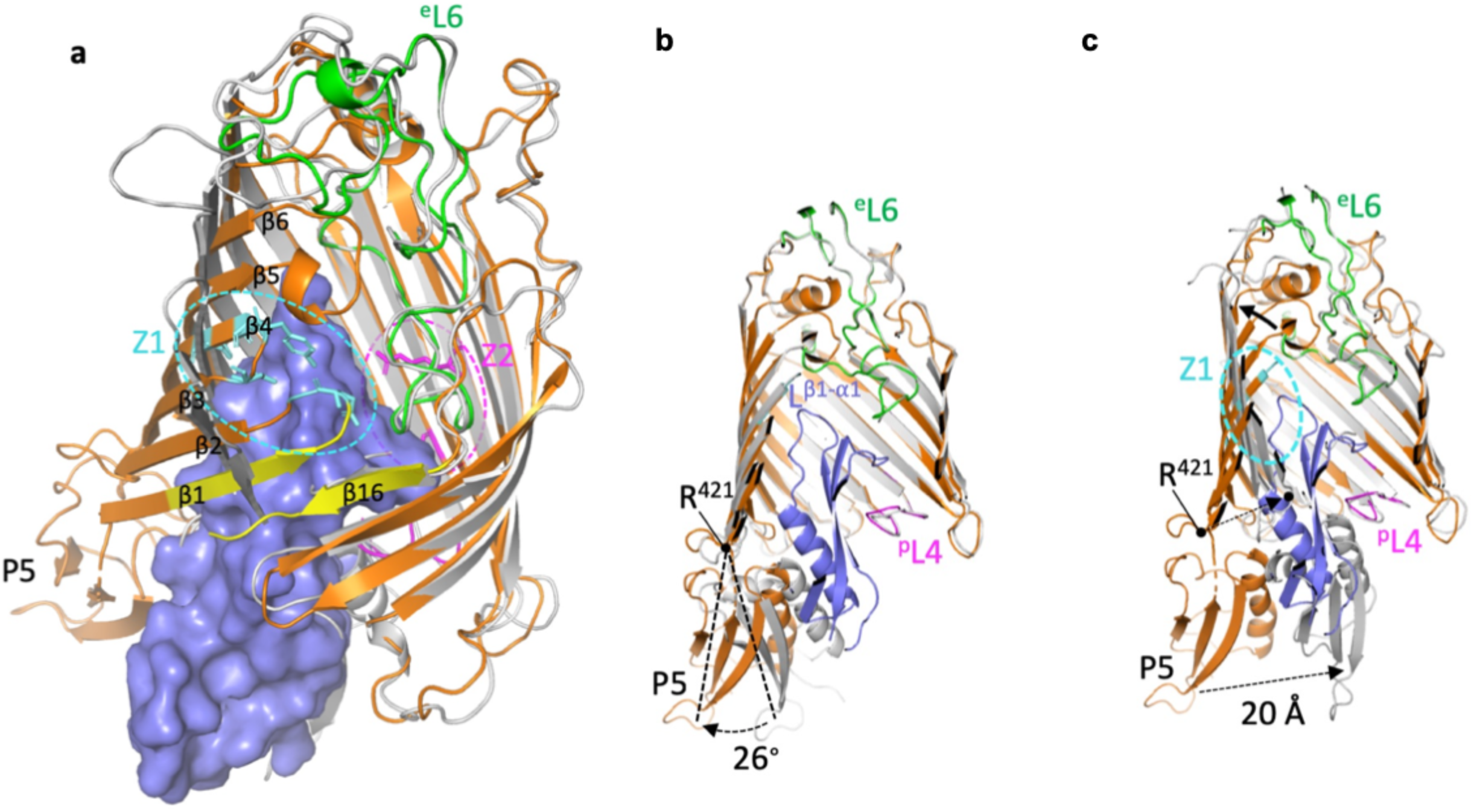
Conformational characteristics of the BamA-RcsF complex. **(a, c)** Tilted top view and slabbed side view of the overlay of the BamA-RcsF complex and BamA in the outward-open conformation (grey, taken from BamACDE complex PDB:5EKQ ^4^). The BamA β-barrel undergoes a ∼45° outward rotation at strands β1-β6, and a 20 Å inward displacement of POTRA5 compared to the structure of BamA-RcsF presented here. **(b)** Slabbed side view of the overlay of BamA-RcsF and BamA in the inward-open conformation (grey, taken from BamABCDE complex PDB:5D0O ^5^). In the structure of BamA-RcsF presented here, POTRA5 makes a 26° outward rotation relative to R421, where it connects to the BamA β-barrel. (a-c) Color scheme for BamA-RcsF is as in Figure 2b. RcsF is shown as a solvent-accessible surface (a) or a ribbon (b, c). Panels (b, c) show side views, slabbed down to view the interior of the complex. For 5EKQ and 5D0O, the Bam accessory proteins BamB, C, D, and E were omitted for clarity, as were POTRA domains 1-4 in all shown BamA structures.

To validate the BamA-RcsF conformation revealed by the X-ray structure, we subjected the complex to crosslinking and analysis via mass spectrometry using the homobifunctional NHS-ester crosslinker disuccinimidyl dibutyric urea ^23^. Crosslinks were identified between lysine residues in RcsF (two lysines from the globular domain and one located at the C-terminus of the linker) and those in POTRA4 and POTRA5 (Extended Data Fig. 4a; Extended Data Table 2), providing further support for the architecture of BamA-RcsF determined by crystallography. To confirm that RcsF binds inside the barrel of BamA, we incorporated the photoreactive lysine analog N6-((3-(3-methyl-3H-diazirin-3-yl)propyl)carbamoyl)-L-lysine (DiZPK) ^24^at multiple positions in the BamA β-barrel domain, selecting residues (R583, R592, K598, K610, R632, R634, R661, K808) whose sidechains face the lumen of the barrel (Extended Data Fig. 4a). After exposure to ultraviolet light, RcsF efficiently crosslinked to BamA when DiZPK was incorporated at three of the selected residues (R592, R598, K610) and to a lower extent at residue K808 (Extended Data Fig. 4a, b), confirming that RcsF binds deep inside the barrel. We measured an equilibrium dissociation constant of 350±49 or 420±48 nM, respectively, depending on whether BamA or RcsF was immobilized (Extended Data Fig. 4c, d). Finally, we deleted loop 1, a short, non-essential ^25^ segment located between the first and second β-strands of the barrel (residues 434 to 437; BamA_Δloop1_), in close proximity to RcsF (Fig. 3a; Extended Data Fig. 4a). RcsF could be pulled down with BamA but not with BamA_Δloop1_ (Extended Data Fig. 4e), indicating that deletion of this loop prevents a stable BamA-RcsF interaction. Further, the Rcs signaling cascade, which is turned on when RcsF fails to interact with BamA ^10^, was constitutively induced in Δ*bamA* cells complemented with BamA_Δloop1_ (Extended Data Fig. 4f). In sum, these results provide functional evidence for our structure of BamA-RcsF and confirm the presence of RcsF inside the barrel of BamA.

Strikingly, our structure suggests that RcsF binding is incompatible with the BamA β-barrel residing in the outward-open conformation (Fig. 3a). Confirming this hypothesis, RcsF was found to bind BamA^G433C/N805C^, a mutant in which opening of the lateral gate is prevented by a disulfide bond between β1 and β16 ^26^, but not to BamA^G393C/G584C^, which is locked in the outward-open conformation ^5^ (Extended Data Fig. 5a, b). However, when reduced, BamA^G393C/G584C^ returned to the inward-open conformation and regained the ability to bind RcsF (Extended Data Fig. 5c). Importantly, given its ability to only bind the inward-open conformation of BamA, the BamA-RcsF complex serves as a proxy for this state. Interestingly, RcsF was recently reported to accumulate on BamA and to jam OMP assembly in the absence of BamB and BamE ^12,13^. Thus, in light of our structural findings, BamA conformational cycling is likely impaired when BamB and BamE are absent. However, binding of these accessory lipoproteins cannot be sufficient to trigger conformational changes in BamA. The structure of BamABCDE has been solved not only in the outward-open conformation but also in the inward-open conformation ^3-6^, despite the presence of all accessory lipoproteins. In addition, we have shown here that RcsF could be co-purified with BamABCDE (Fig. 1; Extended Data Fig. 1), implying that in this purified complex, BamA was in the inward-open conformation. Thus, BamA can remain in the inward-open conformation even when BamBCDE are present, strongly supporting the notion that BAM conformational cycling is triggered by an external signal.

What is this trigger? Insights came from *in vivo* crosslinking experiments carried out in cells in which the expression levels of the BAM components were only moderately increased (∼2-fold) compared with wild-type levels. Whereas RcsF can be crosslinked to BamA when the BamA and BamB subunits are over-expressed, the BamA-RcsF complex becomes barely detectable when all BAM components are moderately overexpressed (Fig. 4a). This observation suggests that the excess copies of BamA-BamB are not functional and do not funnel RcsF to its OMP partners. As a result, BamA-RcsF accumulates and OmpA-RcsF does not form (Fig. 4a). However, if all BAM components (BamABCDE) are moderately overexpressed, normal BAM activity is restored, RcsF only transiently interacts with BamA, and formation of OmpA-RcsF resumes (Fig. 4a). Therefore, whether a stable RcsF complex forms with BAM depends critically on the rates of OMP synthesis and delivery to BAM, as well as the ratio of active BAM complexes to the concentrations of OMP and RcsF substrates. Our data thus support a model in which it is the flux of incoming OMP substrates that triggers conformational changes in the BamA barrel and release of RcsF to its OMP partners (see below; Fig. 4b). Although complexes have to date only been observed between RcsF and OmpA/C/F^10,11^, complexes may form between RcsF and other, less abundant OMPs, depending on the unfolded OMP that is delivered to the BamABCDE-RcsF complex.

**Figure 4.**
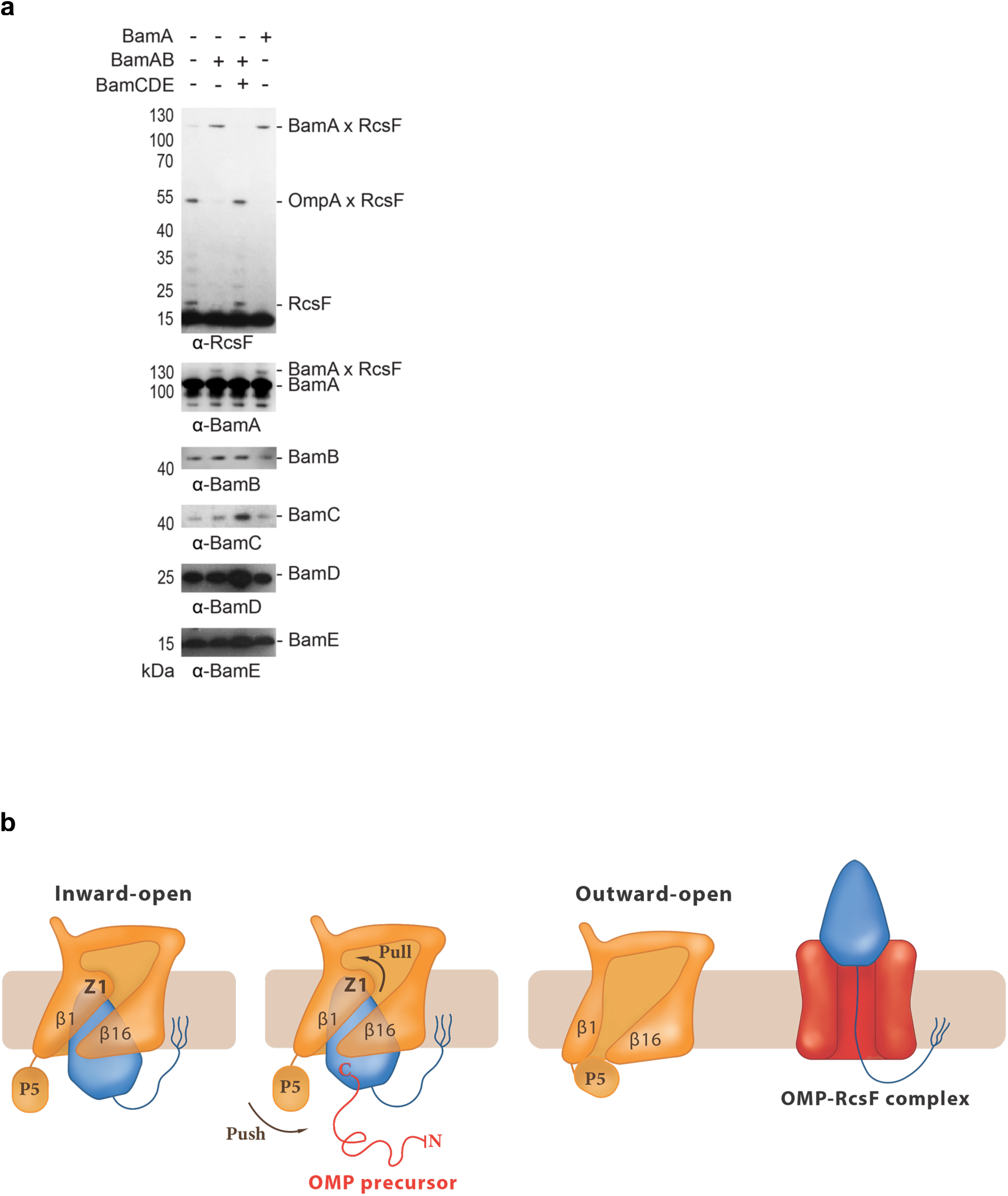
BamA-RcsF is a proxy for the inward-open conformation of BamA. **(a)** *In vivo* chemical crosslinking of RcsF with BamA and OmpA. The BamA-RcsF complex accumulates in cells over-expressing BamA alone or BamA and BamB together. Expression of BamCDE in these cells restores wild-type BamA-RcsF levels. Levels of OmpA-RcsF are inversely correlated with BamA-RcsF. (**b**) Model proposing that BamA conformational cycling is triggered by incoming OMP substrates on the BAM holocomplex. A BamA inward-to-outward open transition could result in an upward displacement of RcsF via a push-and-pull mechanism, resulting in an OMP-RcsF complex. The push-and-pull mechanism involves BamA POTRA5 (P5) and Z1. The topology of the OMP-RcsF complex remains to be established. For clarity, POTRA1-4 and the BAM lipoproteins have been ommitted.

RcsF is not an integral component of BAM; it can bind BamA with high affinity, but it is eventually funneled to OMPs and displayed on the cell surface ^10,11^. It has been proposed that RcsF crosses the outer membrane by being threaded through the lumens of OMPs ^11^. In one possible scenario, RcsF could be transferred from BamA to its OMP partner following opening of the lateral gate and formation of a hybrid barrel (or other non-covalent complex ^9^ between BamA and the nascent OMP), which then buds away from BamA, taking RcsF with it. Available structures show the transition from the inward-open to the outward-open conformation of BamA encompasses a large outward rotation of strands 1-6 of the BamA β-barrel, as well as a 20 Å inward displacement of POTRA 5 (Fig. 3a, b, c) ^5,6^. Strikingly, BamA strands 1-6 coincide with the main RcsF-BamA interaction zone (Z1) seen in our structure, such that outward rotation of Z1 may exert a pulling force on the tip of RcsF (Fig. 3a, c). Concomitantly, the inward movement of POTRA5 would exert a pushing force on the bottom of RcsF (Fig. 3c). We therefore hypothesize that during the inward-to-outward transition of BamA, this push-and-pull action on RcsF could play a role in the transfer of RcsF to its OMP partners and its translocation to the cell surface (Fig. 4b).

It has been proposed that the lipid moiety of RcsF is anchored in the outer leaflet of the membrane and that the N-terminal disordered linker is exposed on the cell surface before being threaded through the lumens of the OMPs ^11^. In this model, the globular domain of RscF resides inside the periplasm. Although we cannot rule out that RcsF flips during release from BamA and transfer to its OMP partners, our structure is more consistent with the hypothesis that BamA releases the globular domain of RcsF on the cell surface. Further investigation will be needed to answer this question, and whether the location of RcsF depends on the identity of its OMP partner.

We previously reported that RcsF uses its interaction with BamA to detect stress in the cell envelope: when damage occurs in the peptidoglycan or the outer membrane, newly synthesized RcsF molecules fail to interact with BamA, activating the Rcs stress response ^10^. Our structure provides a possible explanation for this scenario by suggesting that BamA preferentially adopts the outward-open conformation when envelope integrity is impaired, which would de facto prevent RcsF binding and promote Rcs activation. Thus, we propose that cells could monitor envelope integrity via the conformational cycling of BamA.

By revealing how BamA interacts with RcsF, our work provides insights into the mechanism used by BAM to assemble RcsF-OMP complexes, a novel activity by which BAM exports this lipoprotein to the cell surface. It would be surprising if an essential machinery such as BAM— with a global role in formation of the cell envelope—was only dedicated to export RcsF to the surface. Hence, it is tempting to speculate that other lipoproteins may follow the same route and decorate the surface of *E. coli*, in contrast to the general view that outer membrane lipoproteins face the periplasm ^27^. By showing that the globular domain of RcsF is lodged deep inside the barrel of BamA, our structure also reveals the remarkable—and unanticipated — finding that the BamA β-barrel can accommodate a lipoprotein “substrate” with a globular domain 12 kDa in size. This finding further establishes BAM as an essential hub that contributes to outer membrane biogenesis by interacting both with nascent OMPs for assembly and lipoproteins for export. Future work will reveal whether other lipoproteins bind BamA in a way similar to RcsF. It is remarkable that RcsF binds the lateral gate area and the outward rotating region of the BamA barrel, sites that sense BAM conformational cycling triggered by incoming OMP substrates. Because BAM activity is required for bacterial survival, the complex is an attractive target for new antibiotics. Our work paves the way to the design of new antibacterials that interfere with BAM conformational cycling, because blocking BAM in the inward-open conformation lethally jams BAM with RcsF.

## ACKNOWLEDGMENTS

We thank Asma Boujtat for technical help. We are indebted to Dr. Peng R. Chen for sharing DiZPK, to Dr. Harris Bernstein (NIH, Bethesda, USA) for providing strains and plasmids, and to Dr. Michael Deghelt, Dr. Géraldine Laloux, Dr. Camille Goemans (EMBL, Heidelberg, Germany), and Dr. Pauline Leverrier for helpful suggestions and discussions and for providing comments on the manuscript. We thank Pierre Legrand and staff at Soleil Synchrotron France and at Diamond Light Source UK for beamtime and their assistance during data collection. This work was supported, in part, by grants from the Fonds de la Recherche Scientifique – FNRS, from the FRFS-WELBIO grants n° WELBIO-CR-2015A-03 and WELBIO-CR-2019C-03, from the EOS Excellence in Research Program of the FWO and FRS-FNRS (G0G0818N), from the Fédération Wallonie-Bruxelles (ARC 17/22-087), from the European Commission via the International Training Network Train2Target (721484), and from the BBSRC (BB/P000037/1, BB/M012573/1).

## AUTHOR CONTRIBUTIONS

J.-F.C., R.R.A., H.R., and S.H.C. wrote the manuscript. J.L., RRA, S.N., G.L., S.H.C., H.R., and J.-F.C. designed the experiments. J.L., R.R.A., S.N., G.L., and S.H.C. performed the experiments, constructed the strains, and cloned the constructs. J.L., R.R.A., S.N., S.H.C., H.R., and J.-F.C. analyzed and interpreted the data. A.N.C. and S.E.R. performed and analyzed the crosslinking mass spectrometry experiments. All authors discussed the results and commented on the manuscript.

## AUTHOR INFORMATION

The authors declare no competing financial interests.

## DATA AVAILABILITY

Coordinates and structure factors have been deposited in the Protein Data Bank under accession number 6T1W.

## METHODS

### Bacterial strains, plasmids, and primers

Bacterial strains and plasmids used in this study are listed in Extended Data Tables 3 and 4, respectively. The parental *E. coli* strain DH300 is a MG1655 derivative deleted for the *lac* region and carrying a chromosomal *rprA*_P_*∷lacZ* fusion at the λ phage attachment site to monitor Rcs activation ^28^. To delete *bamA* on the chromosome, a kanamycin resistance (*kan*) cassette ^29^ with the flanking regions of *bamA* was PCR amplified from the genomic DNA of a Δ*rcsF∷kan* strain (PL339) using primers “bamA Km del F” and “bamA Km del R”. Then we performed λ-Red recombineering ^30^ with plasmid pSIM5-tet ^31^ on the strain containing pSC270 as a *bamA*-complementing plasmid in DH300. Deletion of *bamA* was verified by PCR. After preparing P1 lysate from this strain, *bamA* deletion (by transferring the *kan* cassette) was performed via P1 phage transduction of the appropriate strains.

We performed site-directed mutagenesis to generate *bamA* variants on plasmids. For single-codon changes, primer sequences are available upon request; otherwise see Extended Data Table 5. By using pJH114 as a template and performing site-directed mutagenesis (SDM), we introduced a six-histidine tag at the N-terminus of BamA. The C-terminal His-tag of BamE was also removed via SDM, generating pRRA1. The primer pairs were “SDM-HisBamA F” with “SDM-HisBamA R” and “bamE delHis F” with “bamE delHis R”. To add *bamB* next to *bamA* in pBamA, both *bamA* and *bamB* were PCR amplified as a single DNA fragment from pJH114 ^32^ using primers “pTrc-for” and “bamB (NotI) R”. The PCR product and pBamA were digested with NcoI and NotI and then ligated, yielding pBamA-B. To clone *bamC, bamD*, and *bamE* as an operon into a low-copy plasmid (pAM238), PCR was performed on pRA1 as a template using primers “bamC kpnI F” and “pTrc-rev2”. The PCR product and pAM238 were digested with KpnI and XbaI and ligated, generating pSC263. We cloned *bamA* without the six-histidine tag into the low-copy plasmid pSC231^10^, yielding pSC270. *bamA* was PCR amplified from *E. coli* genomic DNA using primers “BamA (PciI)F” and “BamA (XbaI)R” and ligated with pSC231 predigested with NcoI and XbaI. To generate the *bamA* variants locked in the closed and open conformations, SDM was performed on pBamA-B. First, the two cysteines in the ^e^L6 loop of BamA were mutated to serines, generating pBamA_L6_-B. This plasmid was used as template for SDM to generate pBamA^G393C/G584C^-B and pBamA^G433C/N805C^-B. To generate *bamA* variants with amber codons (TAG) to insert 3-(3-methyl-3H-diazirine-3-yl)-propaminocarbonyl-Nε-L-lysine (DiZPK; Artis Chemistry, Shanghai) at selected positions, we performed SDM on pBamA-B and pSC270; primer sequences are available upon request.

### Expression and purification of the BamAB-RcsF complex

*E. coli* PL358 cells harboring pJH118 expressing N-terminal six-histidine-tagged BamA and BamB ^32^ and pSC216 expressing RcsF ^10^ were cultured to overexpress BamA, BamB, and RcsF. Cells were grown in Terrific Broth Auto Inducing Medium (Formedium) supplemented with 0.2% (w/v) L-arabinose at 37 °C (to induce RcsF), ampicillin (200 μg/mL), and chloramphenicol (25 μg/mL). Cells (1 L) were pelleted when they reached OD_600_ ∼ 4, re-suspended in cold phosphate-buffered saline (25 mL) containing a protease-inhibitor cocktail (Complete, Roche), and lysed by two passages through a French pressure cell at 1,500 psi. The cell lysate was centrifuged for 40 min at 40,000 × *g* and 4 °C. After centrifugation, inner-membrane proteins were solubilized using 0.5% (w/v) N-lauryl sarcosine (Sigma) in a buffer containing 20 mM Tris-HCl [pH 7.5] and 150 Mm NaCl for 1.5 h at 4 °C on a roller. The suspension was centrifuged for 40 min at 40,000 x *g* and 4 °C, after which the inner membrane fraction was in the supernatant while the outer membrane fraction remained in the pellet. Outer-membrane proteins were solubilized using 1% (w/v) *n*-dodecyl-β-d-maltopyranoside (DDM; Anatrace) in a buffer containing 20 mM Tris-HCl [pH 7.5], 300 mM NaCl, and 20 mM imidazole overnight at 4 °C on a roller. After centrifugation (40,000 x *g*, 4 °C, 40 min), the supernatant was mixed with Ni-NTA agarose beads (2 mL; IBA Lifescience) equilibrated with 20 mM Tris-HCl [pH 7.5], 300 mM NaCl, 20 mM imidazole, and 1% (w/v) DDM. After washing the resin with 10 column volumes of buffer (20 mM Tris-HCl [pH 7.5], 150 mM NaCl, 20 mM imidazole, 0.6% (w/v) tetraethylene glycol monooctyl ether (C8E4; Anatrace), and 0.01% (w/v) DDM), proteins were eluted with 5 column volumes of the same buffer supplemented with 200 mM imidazole. The eluted complex was then concentrated to 1 mL using a Vivaspin 4 Turbo concentrator (Cut-off 5 kDa; Sartorius). A final purification step was performed using size-exclusion chromatography by loading the proteins on a HiLoad 10/300 Superdex 200 column (GE Healthcare) using 20 mM Tris-HCl [pH 7.5], 150 mM NaCl, 0.6% (w/v) C8E4, and 0.01% (w/v) DDM. Peak fractions were pooled and concentrated to ∼30 mg/mL for crystallization.

For co-crystallization with NaI, NaI replaced NaCl in the gel-filtration buffer. Peak fractions were pooled and concentrated to ∼30 mg/mL using a Vivaspin 4 Turbo concentrator (Sartorius).

### Expression and purification of BAM (BamABCDE) in complex with RcsF

*E. coli* BL21 (DE3) was transformed with pRRA1 expressing all five BAM proteins (N-terminal six-histidine-tagged BamA, BamB, BamC, BamD, and BamE) and pSC216 expressing RcsF for BAM and RcsF overexpression. In control cells, only pRRA1 was transformed. Protein expression and purification were performed as described above except that the detergent was exchanged to 0.1% (w/v) DDM during Ni-NTA affinity chromatography and size-exclusion chromatography. Eluted complexes were identified via SDS-PAGE and concentrated to 4 mg/mL. Blue native electrophoresis of the concentrated complexes was carried out on a 3-12% Bis-Tris gel (Life Technologies) following the manufacturer’s instructions. The protein complex bands separated in the native electrophoresis were identified via SDS-PAGE. Briefly, bands of interest were excised, boiled in SDS-PAGE sample buffer, and applied to the top of a polyacrylamide gel.

### Crystallization, data collection and structure determination

Crystallization assays were carried out using the hanging drop vapor diffusion method in 48-well plates (Molecular Dimensions) at 20°C. The protein solution was mixed in a 2:1 ratio with the crystallization solution from the reservoir. The best native crystals were grown after 4 to 5 days in C10 and G10 conditions from Morpheus crystallization screen (Molecular dimensions; C10: 0.03 M sodium nitrate, 0.03 M sodium phosphate dibasic, 0.03 M ammonium sulfate, 0.10 M Tris-base [pH 8.5]; BICINE, 20 % (v/v) ethylene glycol; 10 % w/v PEG 8000; G10: 0.02 M sodium formate; 0.02 M ammonium acetate; 0.02 M sodium citrate tribasic dihydrate; 0.02M potassium sodium tartrate tetrahydrate; 0.02 M sodium oxamate; 0.10 M Tris-base [pH 8.5]; BICINE; 20% (v/v) ethylene glycol; 10 % (w/v) PEG 8000).

The crystals were harvested in a nylon loop, flash-cooled and stored in liquid nitrogen for data collection. Crystals were screened on beamlines Proxima-1 and Proxima-2 at Synchrotron Soleil (Gif-sur-Yvettes, France) as well as beamlines I03 and I04-1 at Diamond Light Source (Didcot, UK). Data for structure determination were collected on the Proxima-2 beam-line at Synchrotron Soleil at a wavelength of 1.77 Å. Data were indexed and integrated using XDS ^33^, scaled using XSCALE ^33^ and anisotropically corrected using STARANISO, applying a high resolution cutoff of I/σI = 1.2 ^34^. The crystals belong to space group C2, with the unit cell dimensions a=158.84, b=142.5300, c=116.4200 Å^3^ and β=102.61°. The structure was determined by molecular replacement using Phaser ^35^, with the globular domain of RcsF (residues 51-130, PDB:2Y1B) and the inward open BamA β-barrel (residues 422-809, PDB: 5D0O) and BamA POTRA domain 4 (residues 265-344; PDB: 5D0O) as search models. Molecular replacement searches identified two copies each of the BamA β-barrel, RcsF and POTRA domain 4. Following 10 cycles of rigid body refinement POTRA domains 1, 2, 3 and 5 of the first BamA-RcsF copy in the asymmetric unit (Extended Data Figure 2) could be manually placed in the 2FoFc and FoFc difference density and were subjected to an additional 10 rounds of rigid body refinement. The model was refined to 3.8 Å resolution using BUSTER ^36^ and intermittent manual inspection and correction of the model in Coot ^37^. BUSTER was run using Local Structure Similarity Restraints (LSSR) over the two copies in the asymmetric unit, as well as target-based similarity restraints using the inward open BamA structure as reported in PDB:5D0O. The final model shows R and freeR factors of 28.3% and 31.4%, respectively, containing 1388 amino acids, of which 19 are indicated as Ramachandran outliers (1.4%). We note that side chain positioning is frequently ambiguous at 3.8 Å resolution and should therefore not be over-interpreted by users of the deposited model. Side chains for which no unambiguous electron density was observed were not pruned for ease of model interpretation. Such side chains were included in refinement and manually modelled in there most likely rotamer using Coot. Data collection and refinement statistics are found in Extended Data Table 1.

### Site-specific photo-crosslinking

We used a site-specific photo-crosslinking method described previously ^10^ with some modifications. To incorporate DiZPK into BamA, we used the pSup-Mb-DIZPK-RS plasmid encoding an evolved *Methanosarcina barkeri* pyrrolysyl-tRNA synthetase and an optimized 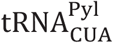 suppressor ^24^. DH300 cells were co-transformed with pSup-Mb-DIZPK-RS and one of the plasmids containing an amber codon in BamA in pSC270. Cells were grown in 3-(N-morpholino) propanesulfonic acid (MOPS) minimal medium supplemented with 0.2% glucose, 0.2% (w/v) arabinose, 200 µM IPTG, 0.001% (w/v) casamino acids, and 0.8 mM DiZPK. When cells reached OD_600_ = 1, 500-µL samples were irradiated with ultraviolet light at 365 nm or left unirradiated for 10 min. Cells were precipitated with trichloroacetic acid, washed with ethanol, and proteins were solubilized in 100 µL SDS-PAGE sample buffer (50 mM Tris-HCl [pH 7.5], 1% (w/v) SDS, 10% (v/v) glycerol, 0.002% (w/v) bromophenol blue) before SDS-PAGE and immunoblotting using anti-RcsF and anti-BamA antibodies.

### *In vivo* BS3 crosslinking

Cells were harvested around mid-log phase (OD_600_ = ∼ 0.5). *In vivo* crosslinking was performed as described previously ^10^, except that bis(sulfosuccinimidyl)suberate (CovaChem) was used instead of 3,3’-dithiobis(sulfosuccinimidyl propionate).

### Chemical crosslinking-mass spectrometry

We first performed buffer exchange of the purified BamAB-RcsF complex using a PD-10 desalting column (GE Healthcare Life Sciences). The complex was eluted with 20 mM HEPES [pH 7.5], 150 mM NaCl, and 0.1% (w/v) DDM. A 30-fold molar excess of the crosslinker disuccinimidyl dibutyric urea (50 mM stock solution in dimethyl sulfoxide, Thermo Scientific) was added to the protein solution and incubated at 37 °C for 1 h. The reaction was quenched by adding Tris-HCl to a final concentration of 20 mM. Crosslinked proteins were precipitated with ethanol and trypsinized, and crosslinked peptides were enriched through cation exchange as described previously ^38^. Briefly, crosslinked proteins (50 µL) were precipitated by adding ice-cold ethanol (150 µL) and 3 M sodium acetate [pH 5.3] (5 µL) prior to incubation at −20 °C for 16 h. The sample was centrifuged (16,200 x *g*, 4 °C, 30 min), the supernatant was removed, and the pellet was washed by adding 80% (v/v) ice-cold ethanol (200 µL) and vortexing for 30 s. The sample was centrifuged again, the supernatant was removed, and the pellet was dried in a vacuum centrifuge. The pellet was dissolved in 1% (w/v) RapiGest (Waters) (10 µL) and trypsin (Sequencing grade, Promega) solution was added (90 µL, 1:50 trypsin:protein mass ratio) before incubating overnight at 37 °C. Trifluoroacetic acid was added (final concentration 0.5% (v/v)) and the sample was incubated at 37 °C for 1 h to precipitate the Rapigest. The mixture was centrifuged (16,200 x *g*, 4 °C, 30 min), the supernatant was concentrated using a vacuum centrifuge, and the pellet was dissolved in 20% (v/v) acetonitrile/0.4% (v/v) formic acid (20 µL). Strong cation exchange enrichment was carried out using OMIX 10 μL strong cation exchange pipette tips (Agilent) as previously described ^38^.

Fractionated peptides (5 µL) were injected onto a reverse-phase Acquity M-Class C18, 75 µm x 150 mm column (Waters) and separated via gradient elution of 1-50% (v/v) solvent B (0.1 % (v/v) formic acid in acetonitrile) in solvent A (0.1 % (v/v) formic acid in water) over 60 min at 300 nL/min. The eluate was infused into a Xevo G2-XS (Waters) mass spectrometer operating in positive ion mode. Mass calibration was performed by infusion of aqueous NaI (2 µg/µL). [Glu1]-Fibrinopeptide B was used for the lock mass spray, with a 0.5 s lock spray scan taken every 30 s. The lock mass correction factor was determined by averaging 10 scans. Data acquisition was performed in DDA mode with a 1 s mass-spectrometry scan over *m*/*z* 350-2000. Instrument parameters were optimized for the detection of crosslinked peptides, as described previously ^39^. Data processing and crosslink identification were performed using MeroX ^40^.

### Expression of BamA mutants and co-purification with RcsF

pBamA and pBamA-B each provide chromosomal-level expression of BamA ^10^. Therefore, we introduced mutations of *bamA* in these plasmids to test the physiological effects of BamA mutants; plasmids were expressed in the presence or absence of *bamA* on the chromosome. Cells (2 mL) were harvested at OD_600_ ∼ 0.5 to purify BamA, except during the following experiment. The cysteine mutants of BamA, when oxidized to form a disulfide bond, allow BamA to form an “open” or “closed” lateral gate. Therefore, the efficiency of disulfide-bond formation in these mutants is very important. To enhance the oxidation of cysteines to form disulfide bonds, we added 3 mM tetrathionate as an oxidant ^41^ at OD_600_ ∼ 0.5 and harvested cells (1 mL) at OD_600_ ∼ 1.0.

Since there was a six-histidine tag at the N-terminus of BamA, we used Dynabeads™ His-Tag (Invitrogen) for Ni-affinity purification. After resuspending cells in 350 µL of 25 mM Tris-HCl [pH 7.4], 290 mM NaCl, 1 mM imidazole, and 0.05% (w/v) DDM (buffer A), cells were lysed via mild sonication on ice. Membrane vesicles were further solubilized by increasing the DDM concentration to 1% (w/v). After removing debris via centrifugation at 9,300 × *g* for 10 min, 5 µL of Dynabeads™ His-Tag (pre-washed with buffer A and resuspended in the same volume) were added to 250 µL of the supernatant, which was incubated for 20 min at 4 °C. The rest of the supernatant was used as the input fraction. The magnetic beads were pulled by a magnet and the supernatant was taken for the flow-through fraction. After washing the beads three times with 750 µL buffer E using the magnet, bound proteins were eluted with 83 µL (three times enrichment compared to the other fractions) of buffer A with 300 mM imidazole. Forty microliters of the input, flow-through, and elution fractions were mixed with SDS-PAGE sample buffer. After denaturation of the three fractions, SDS-PAGE was performed, followed by immunoblotting using rabbit-raised anti-BamA, anti-RcsF ^10^, anti-BamB, anti-BamC, anti-BamD, and anti-BamE.

To determine the redox states of the cysteine-introduced gate mutants of BamA, we added 3 mM N-ethylmaleimide in SDS-PAGE sample buffer to alkylate cysteines to prevent thiol-disulfide exchange. The sample was divided into two aliquots and 10 mM of tris(2-carboxyethyl) phosphine was added to one of them to obtain the reduced state of BamA as a control. Nu-PAGE (4-12% gradient; Novex) was used to separate the oxidized and reduced bands of BamA.

### Biolayer interferometry

Untagged BamA was first biotinylated using the EZ-Link NHS-PEG4-biotin kit (Perbio Science). The reaction was stopped by adding Tris [pH 8] to the final concentration of 20 mM. Excess NHS-PEG4-biotin was removed by passing the sample through a Zeba Spin Desalting column (Perbio Science). Biolayer interferometry was performed in black 96-well plates (Greiner) at 25 °C using OctetRed96 (ForteBio). Streptavidin and Ni-NTA biosensor tips (ForteBio) were hydrated with 0.2 mL working buffer (20 mM Tris [pH 8], 150 mM NaCl, 0.03 % (w/v) DDM) and then loaded with biotinylated BamA or 6xHis-tagged RcsF, respectively.

In the forward experiment, purified 6xHis-tagged RcsF (5 µg/mL) was immobilized on Ni-NTA sensors until the signal reached 0.5-0.6 nm. Association and dissociation of BamA to RcsF-coated tips were monitored for 1200 s and 300 s, respectively, by dipping tips into BamA-containing buffer (serial two-fold dilution from 4000 nM to 62.5 nM), and subsequently in buffer only. In the reverse experiment, biotinylated BamA was immobilized on streptavidin sensor tips to a signal of 2.0 nm. The association and dissociation of RcsF (serial 3-fold dilution from 3000 nM to 12.34 nM) to BamA-coated tips were monitored for 4800 s and 700 s, respectively. Dissociation constants were determined using Graphpad Prism by linear regression of the steady-state binding responses in the saturation binding experiment (Extended Data Fig. 4c, d).

For binding of the BamA^G393C/G584C^ mutant (Extended Data Fig. 5c), 6xHis-tagged RcsF was immobilized on Ni-NTA sensors. To follow BamA association and dissociation, RcsF-coated tips were dipped into 0.2 mL of 200 nM BamA solution, with or without 2 mM diothiothreitol, for 1200 s, followed by 1200 s in buffer only.

### Antibodies and immunoblotting

Rabbit anti-RcsF antibody was previously generated and used by us ^10^. We newly raised the antibodies against BamA, BamB, BamC, BamD, and BamE in rabbits as follows. Except BamA, the DNA sequences encoding the proteins without the signal sequence were cloned into pET28a (Novagen) using the NcoI and XhoI restriction sites, which allows the expressed proteins to be his-tagged at the C-terminus. For BamA, DNA encoding the POTRA domains (1-4) of BamA with a C-terminal strep-tag (but without the signal sequence) was cloned in pET21a (Novagen). All the proteins above were overexpressed in BL21(DE3) and purified using standard methods for Ni-NTA affinity purification or streptavidin purification (POTRA 1-4). Small aliquots of the purified proteins were sent to the CER group (Marloie, Belgium) to raise antibodies in rabbits.

Antibody specificity was confirmed by comparing the immunoblot of the wild-type strain with that of a mutant using each corresponding antibody. The dilutions of the antibodies for immunoblotting were 1:10,000 (BamA), 1:20,000 (BamB), 1:40.000 (BamC), 1:10,000 (BamD), and 1:20,000 (BamE). The specificity of the antibodies was verified; data are available upon request.

To simplify the detection of Bam components and RcsF after purification of BamA, we used two mixtures of antibodies (anti-BamA plus anti-RcsF; anti-BamB, anti-BamC, anti-BamD, plus anti-BamE). Detection specificity was verified using similar mutants as above but harboring pBamA. Data are available upon request.

### β-galactosidase assay

Rcs induction was monitored by measuring β-galactosidase activity as described ^42^.

## LEGENDS TO EXTENDED DATA FIGURES

**Extended Data Figure 1.**
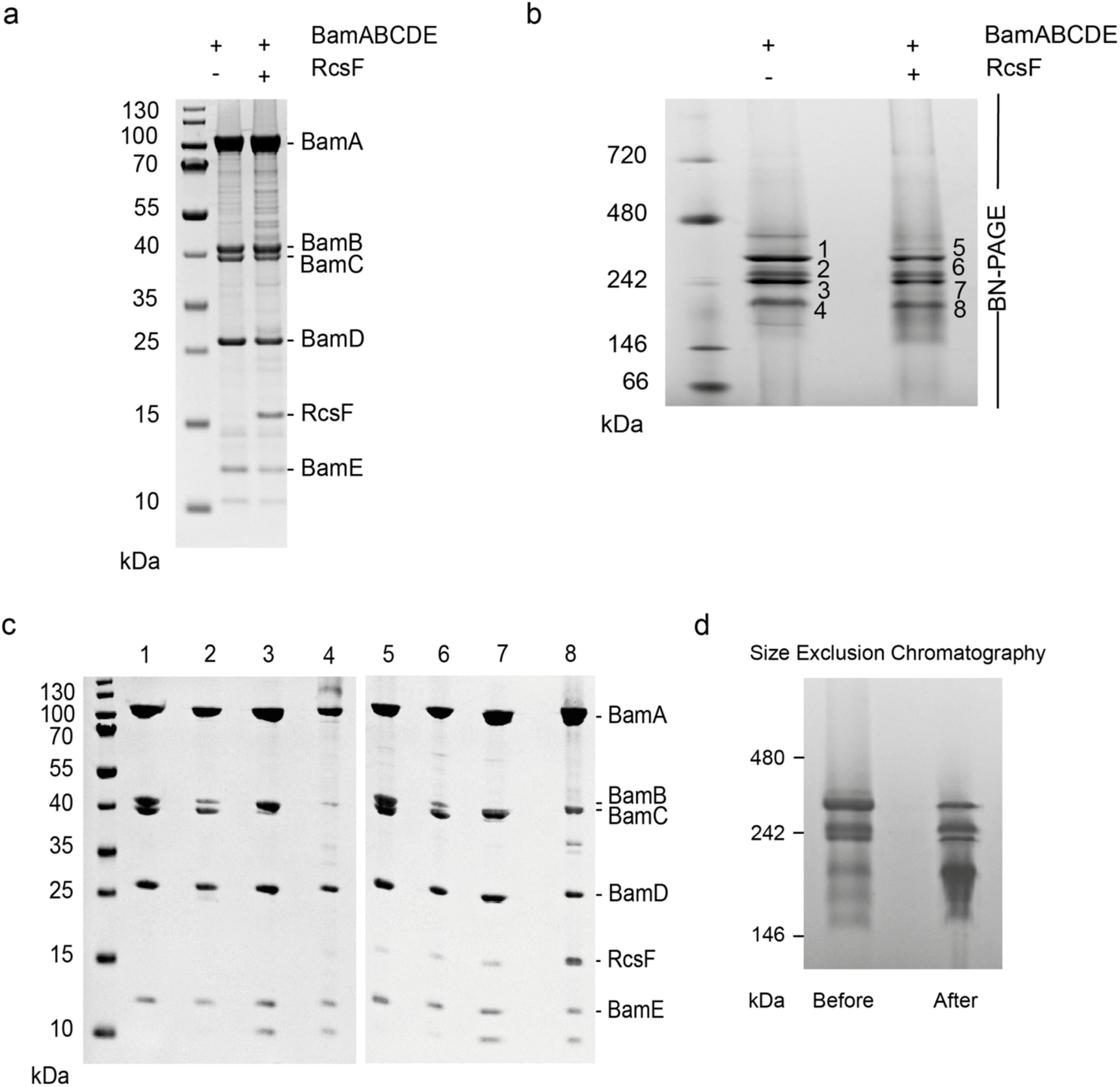
RcsF can be co-purified with the BAM complex. **(a**,**b)** SDS-PAGE (a) and blue native (b) analysis of purified BAM and BAM-RcsF complexes obtained via affinity chromatography. **(c)** SDS-PAGE of the complexes in panel b. The BAM complex expressed from pRRA1 is a mixture of BamABCDE, BamABDE, and BamACDE. When overexpressed, RcsF interacts with all of the complexes. **(d)** Blue native analysis of purified BamABCDE-RcsF before and after size exclusion chromatography, which modifies the migration pattern of the complex, indicating that it is destabilized.

**Extended Data Figure 2.**
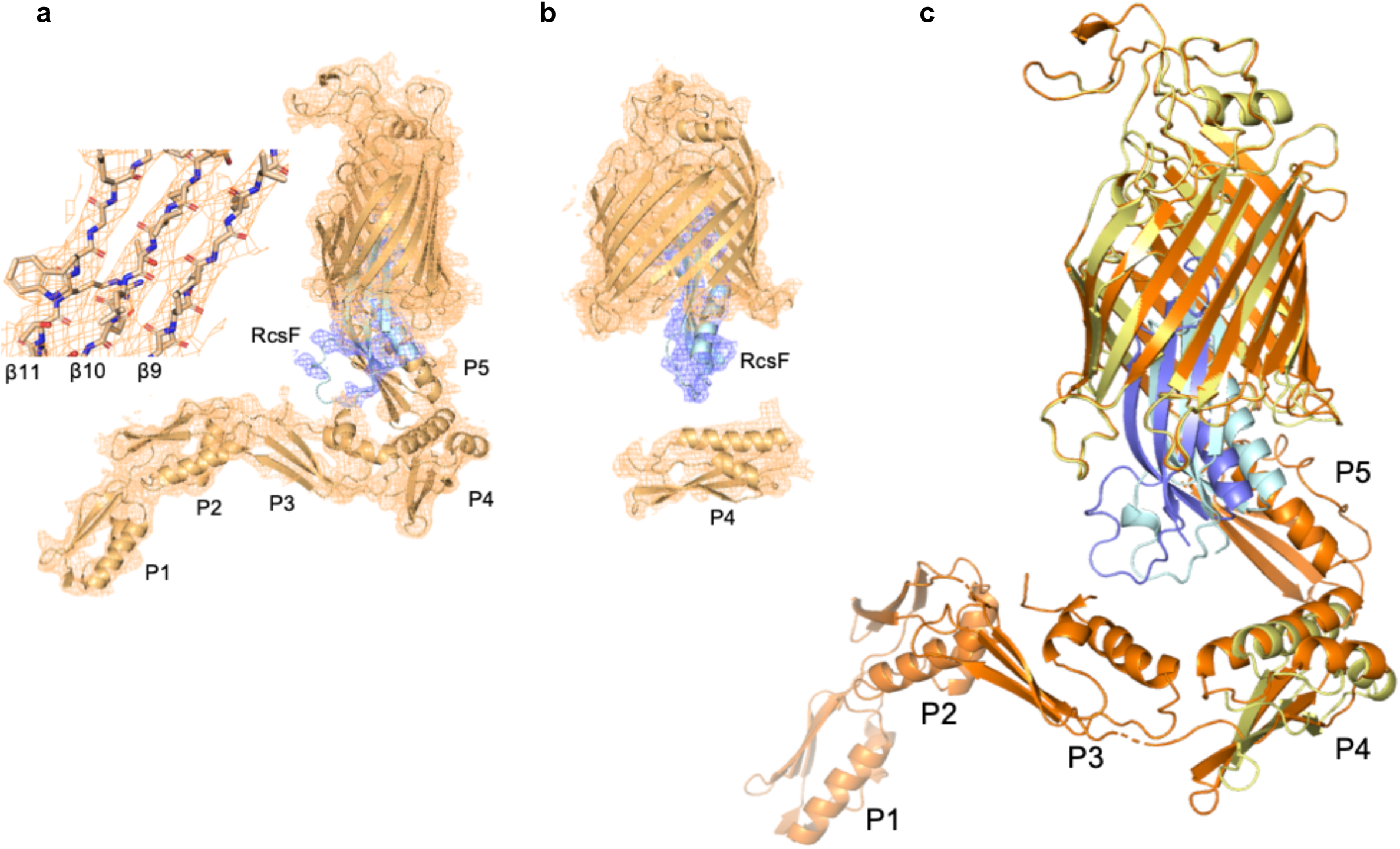
Crystal structure of the BamA-RcsF complex. **(a, b)** Final 2Fo-Fc electron map of the BamA-RcsF complex, shown with a map contour level of 0.08 e-/Å^3^ (root mean square deviation 1.02). The asymmetric unit (AU) of the crystals holds two BamA-RcsF copies, one revealing interpretable electron density for the full BamA sequence (a), and a second revealing unambiguous density for POTRA domain 4 only (b). In the second copy (b), the electron density corresponding to POTRA domains 1, 2, 3, and 5 is too weak to allow unambiguous rigid body placement of the domains. All descriptions and images in the main text are based on the first copy (a). **(c)** Overlay of two BamA-RcsF complexes in the AU. The first complex depicts BamA in gold and RcsF in blue, while these molecules are yellow and light blue, respectively, in the second complex. In both copies, RcsF makes an average displacement of 4 Å relative to the BamA β-barrel.

**Extended Data Figure 3.**
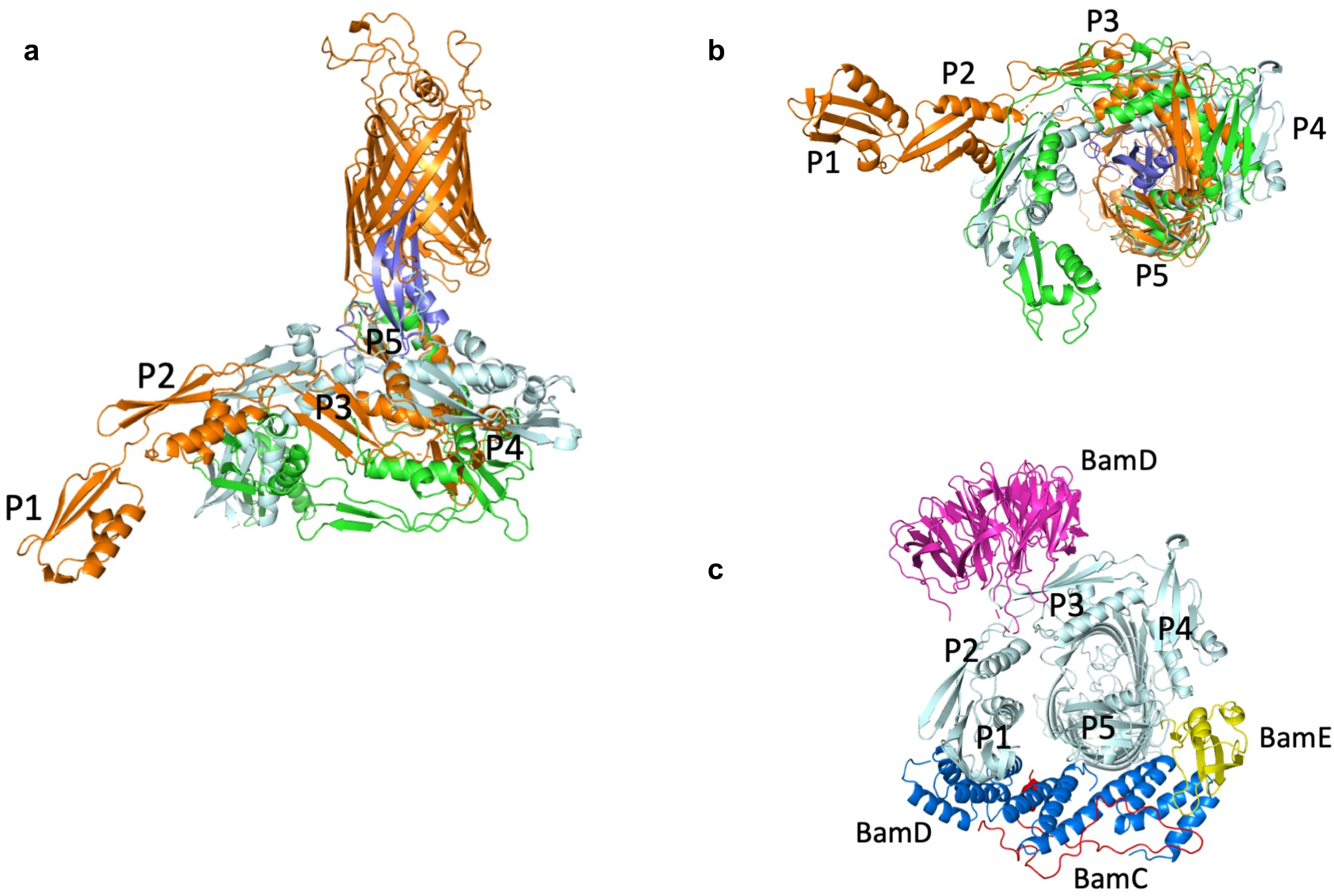
Structural dynamics of the BamA POTRA domains. **(a, b)** Superimposition of BamA-RcsF (gold and blue, respectively) with the POTRA domains in the inward-open BamABCDE complex (PDB: 5D0O; light blue) or the outward-open BamACDE complex (PDB: 5EKQ; green). Complexes are superimposed based on 400 equivalent C*α* atoms in the BamA β-barrel, and shown in side (a) or periplasmic (b) view. For 5d0o and 5ekq, the accessory Bam subunits and the BamA β-barrel are omitted for clarity. **(c)** Periplasmic view of the inward-open BamABCDE complex, showing binding of the Bam accessory proteins BamB (magenta), BamC (red), BamD (blue), and BamE (yellow). Pulldown experiments showed that RcsF binds the BamABCDE complex (Fig. 1). In agreement with this observation, structural comparisons reveal that RcsF binding would not result in direct steric clashes with any Bam accessory protein. However, the positions of the POTRA domains in the BamA-RcsF and BamABCDE complexes are markedly different. In the BamA-RcsF complex, POTRA5 makes a 26° outward rotation to accommodate RcsF (see also Fig. 3), and a reorganization in the joint between POTRA domains 3 and 2 results in a more extended conformation of the POTRA “arm” and the projection of POTRA domains 2 and 1 further from the BamA β-barrel, a conformation not previously reported in available BamA structures. In the BamABCDE complex, BamD contacts both POTRA5 and the joint of POTRA domains 1 and 2. In the BamA-RcsF complex, POTRA5 and POTRA domains 2 and 1 are too distant to be bridged by BamD; binding of BamD to BamA-RcsF therefore requires a conformational change in the POTRA arm or the dissociation of BamD at either of these two contact points.

**Extended Data Figure 4.**
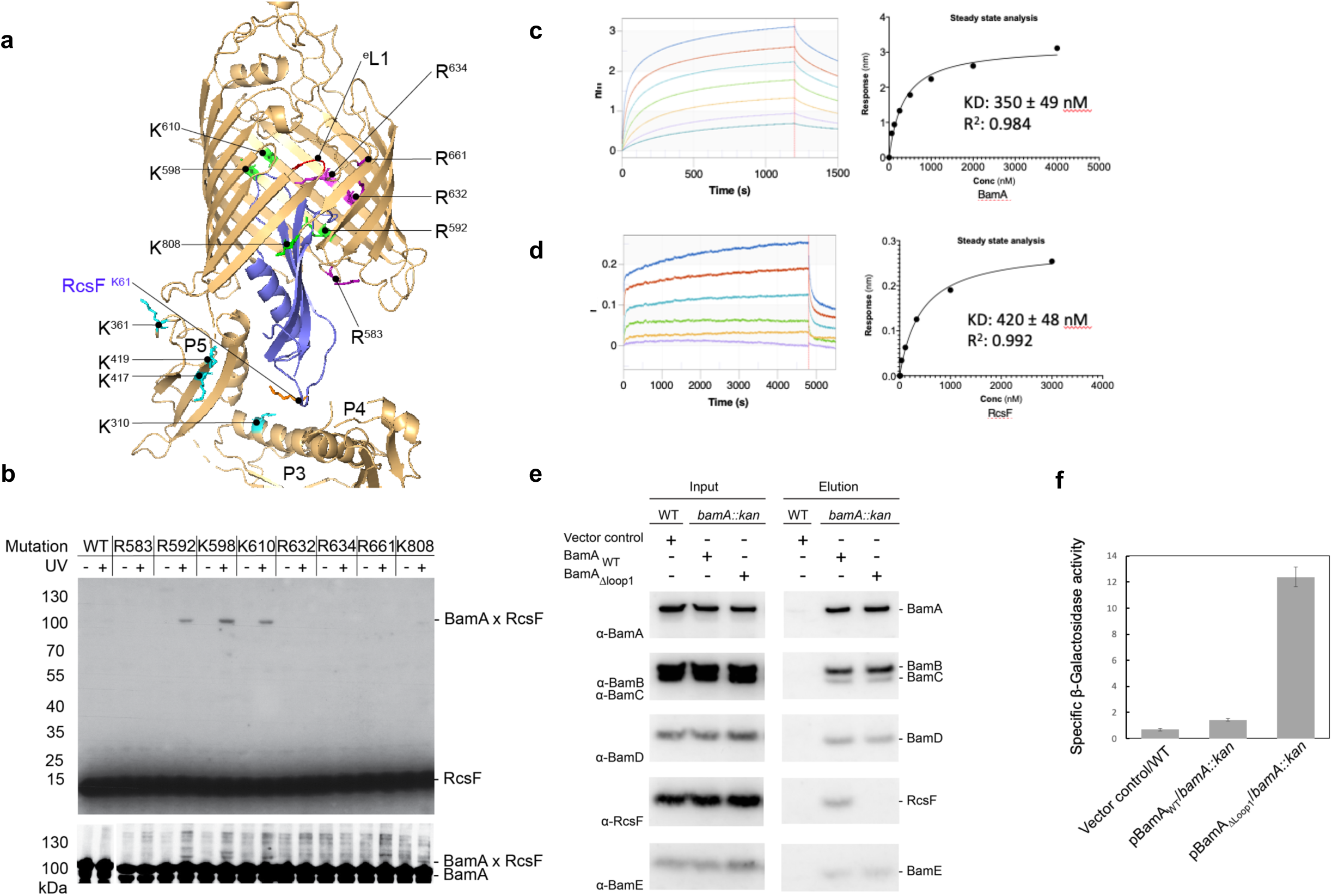
Validation of the BamA-RcsF structure. **(a)** Ribbon diagram of the BamA-RcsF structure. Highlighted residues show sites mutated to amber for incorporation of the photoreactive lysine analog DiZPK. Sites that crosslink to RcsF are green, sites that show no crosslinking are magenta. Mutation of extracellular loop 1 (^e^L1; red) leads to loss of RcsF binding (see panel b). BamA sidechains found to crosslink with RcsF by means of the homobifunctional amine-reactive crosslinker disuccinimidyl dibutyric urea (DSBU) are shown as sticks and colored cyan. Residue K61 from RcsF, which was found to crosslink to BamA using DSBU, is shown as a stick and colored orange. The other two RcsF residues (K42 and K134) that could be crosslinked to BamA are not visible in this structural model. **(b)** *In vivo* photocrosslinking experiment in which cells expressing the BamA mutants containing DiZPK at the indicated positions were treated (+) or not (-) with ultraviolet light. Proteins samples were analyzed via SDS-PAGE and immunoblotted with anti-RcsF and anti-BamA antibodies, showing that the photo-crosslinked complexes contain BamA and RcsF. WT, wild type. **(c**,**d)** Sensorgrams from biolayer interferometry (left) and corresponding equilibrium binding plots (right) of immobilized RcsF titrated with BamA (c) or immobilized BamA titrated with RcsF (d). **(e)** Deletion of loop 1 in BamA prevents RcsF from being pulled down with BamA. WT cells harboring the empty plasmid (pAM238) were used as control. **(f)** Overexpression of pBamA_ΔLoop1_in a *bamA* deletion strain activates the Rcs system. A chromosomal *rprA∷lacZ* fusion was used to monitor Rcs activity, and specific β-galactosidase activity was measured from cells at mid-log phase (OD_600_=0.5). Error bars depict standard deviation (n=4).

**Extended Data Figure 5.**
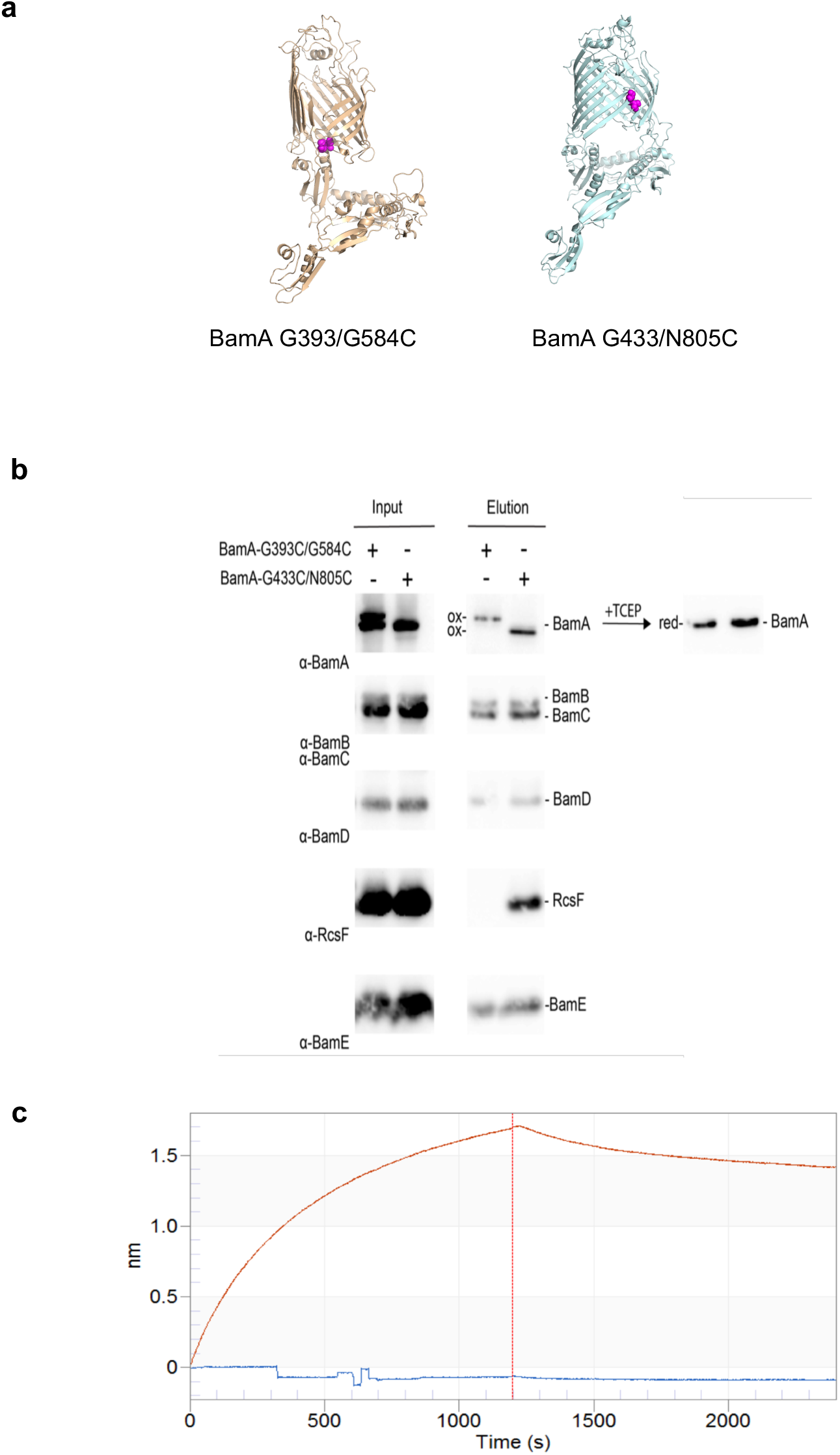
RcsF binds the inward-open conformation of BamA. **(a)** Models for the BamA^G393C/G584C 5^ and BamA^G433C/N805C 26^ double cysteine mutants, which are locked in the outward-open or inward-open conformation, respectively, when oxidized. Mutated cysteines are shown as atom spheres. (**b**) BamA barrel locking and RcsF binding. Overexpression of double cysteine mutants pBamA^G393C/G584C^-B and BamA^G433C/N805C^-B in a wild-type strain. RcsF can be co-purified with the BamA β-barrel locked in the inward-open conformation (BamA^G433C/N805C^) by a disulfide bond (ox) but not in the outward-open conformation (BamA^G393C/G584C^). BamA mutants become reduced (red) following treatment with tris(2-carboxyethyl) phosphine (TCEP) and migrate similarly. (**c**) Sensorgram from biolayer interferometry of immobilized BamA^G393C/G584C^ titrated with RcsF, without (oxidized; -DTT) or with dithiothreitol (reduced; +DTT). When the β-barrel is locked in the outward-open conformation (-DTT), RcsF is unable to bind BamA. When reduced, BamA^G393C/G584C^ regains binding, demonstrating that BamA reverts to the inward-open conformation in which it can bind RcsF.

## EXTENDED DATA TABLES

**Extended Data Table 1.** Data collection and refinement statistics

|  | **BamA-RcsF** |
| --- | --- |
| **Data collection** |  |
| Space group | C2 |
| Cell dimensions |  |
| *a*, *b*, *c* (Å) | 158.84, 142.53, 116.42 |
| $\alpha$, $\beta$, $\gamma$ (°) | 90.0, 102.6, 90.0 |
| Resolution (Å)** | 48.6 – 3.8 (4.0 – 3.8) * |
| *R*<sub>pim</sub> | 0.02 (0.79) |
| *I* / $\sigma I$ | 15.9 (0.8) |
| Completeness (%) |  |
| Spherical | 71.2 (24.9) |
| Elliptical** | 92.8 (91.4) |
| Redundancy | 34.3 (31.7) |
| Wilson B (Å<sup>2</sup>) | 149.4 |
| **Refinement** |  |
| Resolution (Å) | 48.6 – 3.8 |
| No. reflections | 18489 |
| *R*<sub>work</sub> / *R*<sub>free</sub> | 28.1 / 31.7 |
| No. atoms |  |
| Protein | 10838 |
| Ligand/ion | NA |
| Water | NA |
| *B*-factors (Å<sup>2</sup>) |  |
| Protein | 193.5 |
| Ligand/ion | NA |
| Water | NA |
| R.m.s. deviations |  |
| Bond lengths (Å) | 0.01 |
| Bond angles (°) | 1.15 |

\*Values in parentheses are for highest-resolution shell.
\*\* Elliptical diffraction limits and principal reciprocal axes of the fitted ellipsoid: 4.6 Å along 0.987 a\* - 0.158 c\*, 4.4 Å along b\* and 3.8 Å along c\*

**Extended Data Table 2.** Intermolecular BamA-RcsF crosslinks detected. PSMs = Peptide Spectrum Matches. The POTRA domain in BamA where the crosslinked residue is located is also indicated.

| **BamA**<br>**Residue** | **RcsF**<br>**Residue** | **BamA Peptide** |  |  | **RcsF Peptide** |  |  | **PSMs** | **POTRA** |
| --- | --- | --- | --- | --- | --- | --- | --- | --- | --- |
|  |  | **Sequence** | **Start** | **End** | **Sequence** | **Start** | **End** |  |  |
| 310 | 61 | **KLLGR** | 310 | 314 | IYTNAEELVG**K**PFR | 51 | 64 | 3 | 4 |
| 361 | 42 | FEGND**T**SKDAVLR | 354 | 366 | AEPAKP**K**APR | 36 | 45 | 3 | 5 |
| 361 | 61 | FEGND**T**SKDAVLR | 354 | 366 | IYTNAEELVG**K**PFR | 51 | 64 | 78 | 5 |
| 310 | 134 | **KLLGR** | 310 | 314 | QAVCIGSALNIT**A**K | 121 | 134 | 1 | 4 |
| 417 | 61 | VPGSPDQVDVVY**K**VK | 405 | 419 | IYTNAEELVG**K**PFR | 51 | 64 | 14 | 5 |
| 419 | 61 | **V**KER | 418 | 421 | IYTNAEELVG**K**PFR | 51 | 64 | 24 | 5 |

N.B. Residue 134 is the C-terminal Lys of RscF

**Extended Data Table 3.** Strains used in this study.

| Strains | Genotype and description | Source |
| --- | --- | --- |
| BL21<br>(DE3) | F- *ompT hsdSB* (rB- mB-) *gal dcm* (DE3) | Novagen |
| DH300 | *rprA-lacZ* MG1655 (*argF-lac*) *U169* | 28 |
| PL339 | $\Delta rcsF::kan$ | 10 |
| PL358 | DH300 $\Delta rcsF$ | 10 |
| Keio<br>collection<br>single<br>mutants | $\Delta rcsF::kan$ | 43 |
| RRA73 | BL21 (DE3) with pRRA1 | This study |
| RRA74 | BL21 (DE3) with pRRA1 and pSC216 | This study |
| SEN1071 | PL358 with pJH118 and pSC216 | This study |
| SEN1472 | DH300 $\Delta bamA::kan$ with pSC270 | This study |
| SEN1603 | DH300 $\Delta bamA::kan$ with pBamA | This study |
| SEN1607 | DH300 $\Delta bamA::kan$ with pBamAΔloop1 | This study |
| SEN1411 | DH300 with pET3a and pAM238 | This study |
| SEN991 | DH300 with pBamA and pAM238 | This study |
| SEN1597 | DH300 with pBamA-B and pAM238 | This study |
| SEN1598 | DH300 with pBamA-B and pSC263 | This study |
| RRA104 | DH300 with pBamA<sub>L6</sub>-B and pAM238 | This study |
| RRA54 | DH300 with pBamA<sup>G393C/G584C</sup>-B and pAM238 | This study |
| RRA106 | DH300 with pBamA<sup>G433C/N805C</sup>-B and pAM238 | This study |

**Extended Data Table 4.**
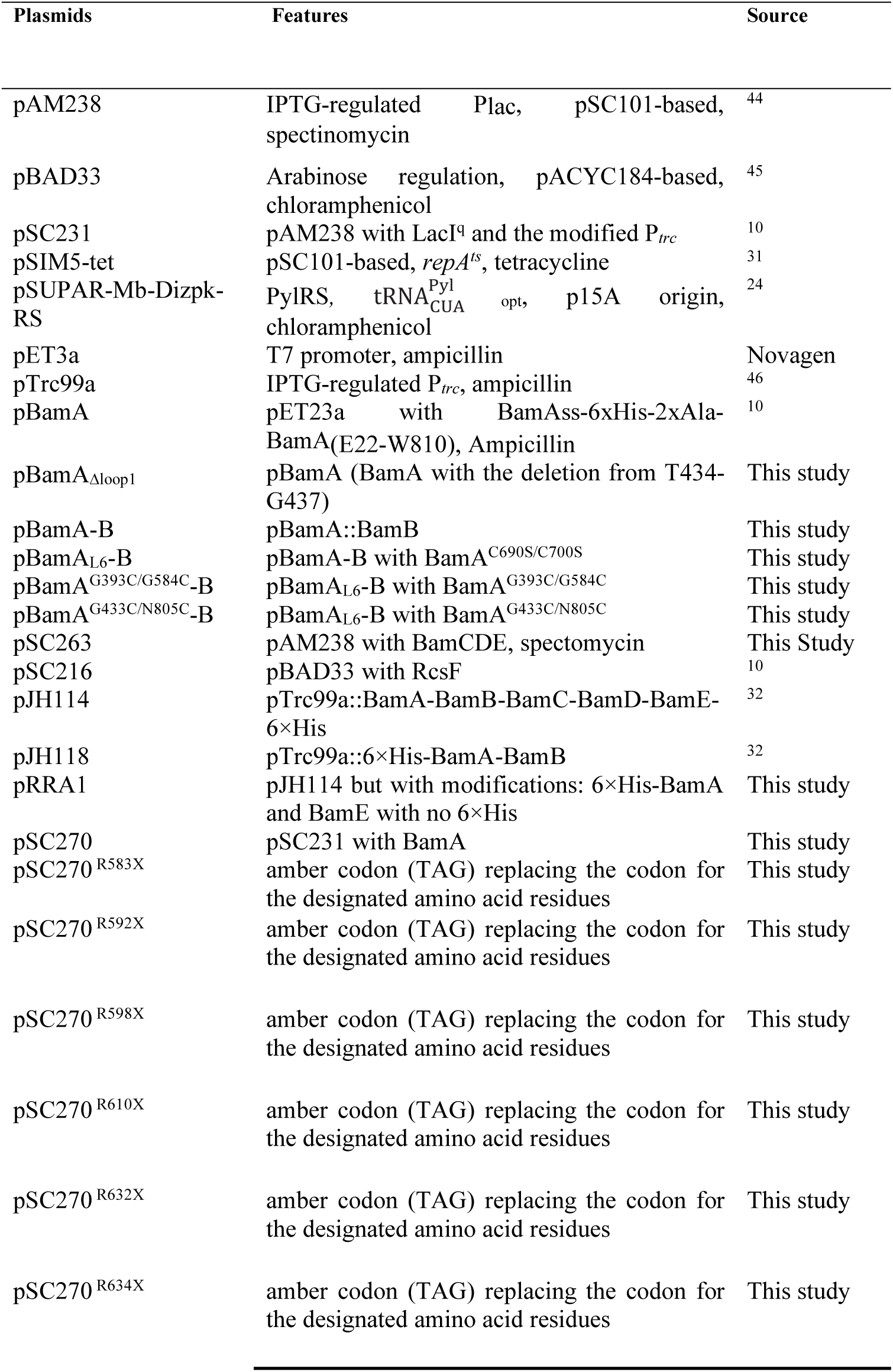
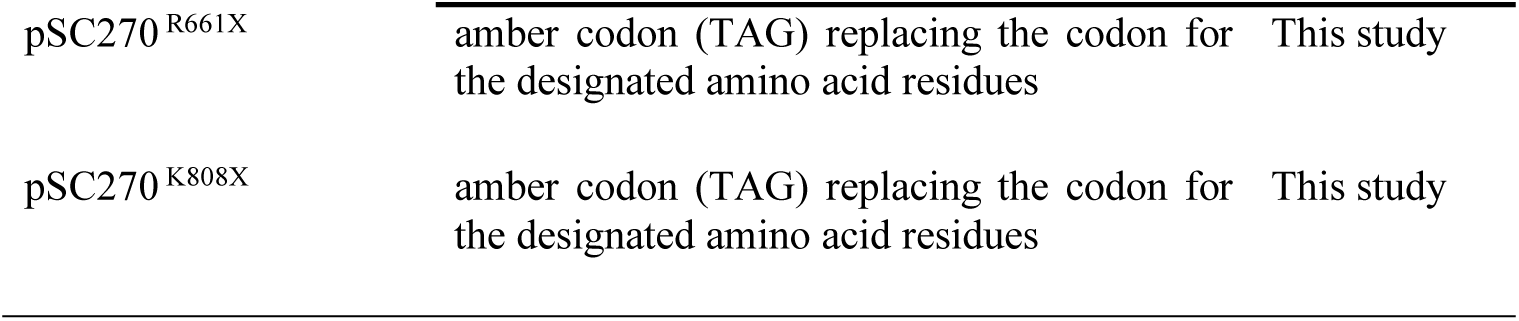
Plasmids used in this study.

**Extended Data Table 5.**
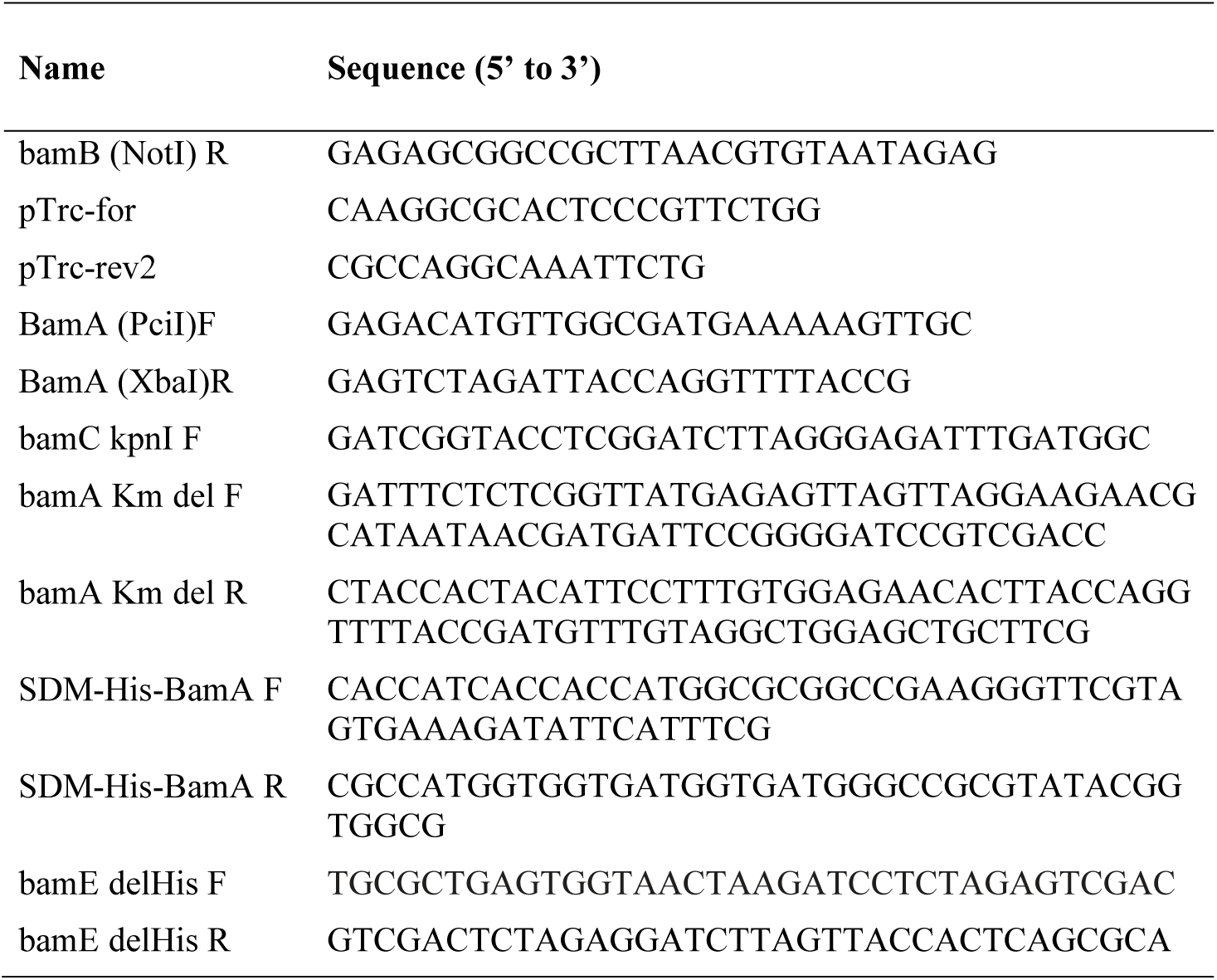
Primers used in this study.

## REFERENCES

1 Noinaj, N., Gumbart, J. C. & Buchanan, S. K. The beta-barrel assembly machinery in motion. Nat Rev Microbiol 15, 197–204, doi:10.1038/nrmicro.2016.191 (2017).

2 Hagan, C. L., Silhavy, T. J. & Kahne, D. beta-Barrel membrane protein assembly by the Bam complex. Annu Rev Biochem 80, 189–210, doi:10.1146/annurev-biochem-061408-144611 (2011).

3 Iadanza, M. G. et al. Lateral opening in the intact beta-barrel assembly machinery captured by cryo-EM. Nat Commun 7, 12865, doi:10.1038/ncomms12865 (2016).

4 Bakelar, J., Buchanan, S. K. & Noinaj, N. The structure of the beta-barrel assembly machinery complex. Science 351, 180–186, doi:10.1126/science.aad3460 (2016).

5 Gu, Y. et al. Structural basis of outer membrane protein insertion by the BAM complex. Nature 531, 64–69, doi:10.1038/nature17199 (2016).

6 Han, L. et al. Structure of the BAM complex and its implications for biogenesis of outer-membrane proteins. Nat Struct Mol Biol 23, 192–196, doi:10.1038/nsmb.3181 (2016).

7 Wu, T. et al. Identification of a multicomponent complex required for outer membrane biogenesis in Escherichia coli. Cell 121, 235–245 (2005).

8 Sklar, J. G. et al. Lipoprotein SmpA is a component of the YaeT complex that assembles outer membrane proteins in Escherichia coli. Proc Natl Acad Sci U S A 104, 6400–6405 (2007).

9 Schiffrin, B., Brockwell, D. J. & Radford, S. E. Outer membrane protein folding from an energy landscape perspective. BMC Biol 15, 123, doi:10.1186/s12915-017-0464-5 (2017).

10 Cho, S. H. et al. Detecting Envelope Stress by Monitoring beta-Barrel Assembly. Cell 159, 1652–1664, doi:10.1016/j.cell.2014.11.045 (2014).

11 Konovalova, A., Perlman, D. H., Cowles, C. E. & Silhavy, T. J. Transmembrane domain of surface-exposed outer membrane lipoprotein RcsF is threaded through the lumen of beta-barrel proteins. Proc Natl Acad Sci U S A 111, E4350–4358, doi:10.1073/pnas.1417138111 (2014).

12 Tata, M. & Konovalova, A. Improper Coordination of BamA and BamD Results in Bam Complex Jamming by a Lipoprotein Substrate. MBio 10, doi:10.1128/mBio.00660-19 (2019).

13 Hart, E. M., Gupta, M., Wuhr, M. & Silhavy, T. J. The Synthetic Phenotype of DeltabamB DeltabamE Double Mutants Results from a Lethal Jamming of the Bam Complex by the Lipoprotein RcsF. MBio 10, doi:10.1128/mBio.00662-19 (2019).

14 Wall, E., Majdalani, N. & Gottesman, S. The Complex Rcs Regulatory Cascade. Annu Rev Microbiol 72, 111–139, doi:10.1146/annurev-micro-090817-062640 (2018).

15 Laloux, G. & Collet, J. F. “Major Tom to ground control: how lipoproteins communicate extra-cytoplasmic stress to the decision center of the cell". J Bacteriol, doi:10.1128/JB.00216-17 (2017).

16 Hussein, N. A., Cho, S. H., Laloux, G., Siam, R. & Collet, J. F. Distinct domains of Escherichia coli IgaA connect envelope stress sensing and down-regulation of the Rcs phosphorelay across subcellular compartments. PLoS Genet 14, e1007398, doi:10.1371/journal.pgen.1007398 (2018).

17 Kaur, H. et al. Identification of conformation-selective nanobodies against the membrane protein insertase BamA by an integrated structural biology approach. J Biomol NMR 73, 375–384, doi:10.1007/s10858-019-00250-8 (2019).

18 Albrecht, R. et al. Structure of BamA, an essential factor in outer membrane protein biogenesis. Acta Crystallogr D Biol Crystallogr 70, 1779–1789, doi:10.1107/S1399004714007482 (2014).

19 Ni, D. et al. Structural and functional analysis of the beta-barrel domain of BamA from Escherichia coli. FASEB J 28, 2677–2685, doi:10.1096/fj.13-248450 (2014).

20 Hartmann, J. B., Zahn, M., Burmann, I. M., Bibow, S. & Hiller, S. Sequence-Specific Solution NMR Assignments of the beta-Barrel Insertase BamA to Monitor Its Conformational Ensemble at the Atomic Level. J Am Chem Soc 140, 11252–11260, doi:10.1021/jacs.8b03220 (2018).

21 Leverrier, P. et al. Crystal structure of the outer membrane protein RcsF, a new substrate for the periplasmic protein-disulfide isomerase DsbC. J Biol Chem 286, 16734–16742, doi:10.1074/jbc.M111.224865 (2011).

22 Rogov, V. V., Rogova, N. Y., Bernhard, F., Lohr, F. & Dotsch, V. A disulfide bridge network within the soluble periplasmic domain determines structure and function of the outer membrane protein RCSF. J Biol Chem 286, 18775–18783, doi:10.1074/jbc.M111.230185 (2011).

23 Calabrese, A. N. & Radford, S. E. Mass spectrometry-enabled structural biology of membrane proteins. Methods 147, 187–205, doi:10.1016/j.ymeth.2018.02.020 (2018).

24 Zhang, M. et al. A genetically incorporated crosslinker reveals chaperone cooperation in acid resistance. Nat Chem Biol 7, 671–677, doi:10.1038/nchembio.644 (2011).

25 Gu, Y., Zeng, Y., Wang, Z. & Dong, C. BamA beta16C strand and periplasmic turns are critical for outer membrane protein insertion and assembly. Biochem J 474, 3951–3961, doi:10.1042/BCJ20170636 (2017).

26 Noinaj, N., Kuszak, A. J., Balusek, C., Gumbart, J. C. & Buchanan, S. K. Lateral opening and exit pore formation are required for BamA function. Structure 22, 1055–1062, doi:10.1016/j.str.2014.05.008 (2014).

27 Okuda, S. & Tokuda, H. Lipoprotein sorting in bacteria. Annu Rev Microbiol 65, 239–259, doi:10.1146/annurev-micro-090110-102859 (2011).

28 Majdalani, N., Hernandez, D. & Gottesman, S. Regulation and mode of action of the second small RNA activator of RpoS translation, RprA. Mol Microbiol 46, 813–826 (2002).

29 Datsenko, K. A. & Wanner, B. L. One-step inactivation of chromosomal genes in Escherichia coli K-12 using PCR products. Proc Natl Acad Sci U S A 97, 6640–6645, doi:10.1073/pnas.120163297120163297 [pii] (2000).

30 Yu, D. et al. An efficient recombination system for chromosome engineering in Escherichia coli. Proc Natl Acad Sci U S A 97, 5978–5983, doi:10.1073/pnas.100127597 (2000).

## REFERENCES

31 Koskiniemi, S., Pranting, M., Gullberg, E., Nasvall, J. & Andersson, D. I. Activation of cryptic aminoglycoside resistance in Salmonella enterica. Mol Microbiol 80, 1464–1478, doi:10.1111/j.1365-2958.2011.07657.x (2011).

32 Roman-Hernandez, G., Peterson, J. H. & Bernstein, H. D. Reconstitution of bacterial autotransporter assembly using purified components. Elife 3, e04234, doi:10.7554/eLife.04234 (2014).

33 Kabsch, W. Xds. Acta Crystallogr. D Biol. Crystallogr. 66, 125–132 (2010).

34 Tickle, I. J., Flensburg, C., Keller, P., Paciorek, W., Sharff, A., Vonrhein, C., Bricogne, G.. STARANISO, 2018).

35 McCoy, A. J. et al. Phaser crystallographic software. J Appl Crystallogr 40, 658–674, doi:10.1107/S0021889807021206 (2007).

36 Bricogne G. B. E., Brandl M., Flensburg C., Keller P., Paciorek W., & Roversi P, S. A., Smart O.S., Vonrhein C., Womack T.O. BUSTED version 2.10.3. (2017).

37 P. Emsley, K. C. Coot: model-building tools for molecular graphics. Acta Crystallogr. D Biol. Crystallogr. 60, 2126–2132 (2004).

38 Schmidt, C. & Robinson, C. V. A comparative cross-linking strategy to probe conformational changes in protein complexes. Nat Protoc 9, 2224–2236, doi:10.1038/nprot.2014.144 (2014).

39 James, J. M. B., Cryar, A. & Thalassinos, K. Optimization Workflow for the Analysis of Cross-Linked Peptides Using a Quadrupole Time-of-Flight Mass Spectrometer. Anal Chem 91, 1808–1814, doi:10.1021/acs.analchem.8b02319 (2019).

40 Iacobucci, C. et al. A cross-linking/mass spectrometry workflow based on MS-cleavable cross-linkers and the MeroX software for studying protein structures and protein-protein interactions. Nat Protoc 13, 2864–2889, doi:10.1038/s41596-018-0068-8 (2018).

41 Osborne, A. R. & Rapoport, T. A. Protein translocation is mediated by oligomers of the SecY complex with one SecY copy forming the channel. Cell 129, 97–110, doi:10.1016/j.cell.2007.02.036 (2007).

42 Miller, J. C. Experiments in Molecular Genetics. (Cold Spring Harbor Laboratory Press, 1972).

43 Baba, T. et al. Construction of Escherichia coli K-12 in-frame, single-gene knockout mutants: the Keio collection. Mol Syst Biol 2, 2006 0008, doi:10.1038/msb4100050 (2006).

44 Gil, D. & Bouche, J. P. ColE1-type vectors with fully repressible replication. Gene 105, 17–22 (1991).

45 Guzman, L. M., Belin, D., Carson, M. J. & Beckwith, J. Tight regulation, modulation, and high-level expression by vectors containing the arabinose PBAD promoter. J Bacteriol 177, 4121–4130 (1995).

46 Weiss, D. S., Chen, J. C., Ghigo, J. M., Boyd, D. & Beckwith, J. Localization of FtsI (PBP3) to the septal ring requires its membrane anchor, the Z ring, FtsA, FtsQ, and FtsL. J Bacteriol 181, 508–520 (1999).

